# Catabolism of 3-hydroxypyridine by *Ensifer adhaerens* HP1: a novel four-component gene encoding 3-hydroxypyridine dehydrogenase HpdA catalyzes the first step of biodegradation

**DOI:** 10.1101/2020.01.08.898148

**Authors:** Haixia Wang, Xiaoyu Wang, Hao Ren, Xuejun Wang, Zhenmei Lu

## Abstract

3-Hydroxypyridine (3HP) is an important natural pyridine derivative. *Ensifer adhaerens* HP1 can utilize 3HP as the sole source of carbon, nitrogen and energy to grow. However, the genes responsible for the degradation of 3HP remain unknown. In this study, we predicted that a gene cluster, designated *3hpd*, may be responsible for the degradation of 3HP. The initial hydroxylation of 3HP is catalyzed by a four-component dehydrogenase (HpdA1A2A3A4), leading to the formation of 2,5-dihydroxypyridine (2,5-DHP) in *E. adhaerens* HP1. In addition, the SRPBCC component in HpdA existed as a separate subunit, which is different from other SRPBCC-containing molybdohydroxylases acting on N-heterocyclic aromatic compounds. Our findings provide a better understanding of the microbial degradation of pyridine derivatives in nature. Additionally, research on the origin of the discovered four-component dehydrogenase with a separate SRPBCC domain may be of great significance.

**Importance:** 3-Hydroxypyridine is an important building block for synthesizing drugs, herbicides and antibiotics. Although the microbial degradation of 3-hydroxypyridine has been studied for many years, the molecular mechanisms remain unclear. Here, we show that *3hpd* is responsible for the catabolism of 3-hydroxypyridine. The *3hpd* gene cluster was found to be widespread in *Actinobacteria*, *Rubrobacteria*, *Thermoleophilia*, and *Alpha-*, *Beta-*, and *Gammaproteobacteria*, and the genetic organization of the *3hpd* gene clusters in these bacteria showed high diversity. Our findings provide new insight into the catabolism of 3-hydroxypyridine in bacteria.

## Introduction

The pyridine ring is a major constituent of natural compounds such as plant alkaloids, coenzymes and antibiotics. 3-Hydroxypyridine (3HP), a useful and valuable pyridine derivative, is a monohydroxypyridine in which the hydrogen at position 3 of the pyridine has been replaced by a hydroxyl group. 3HP has been detected as a thermal degradation product in the smoke from burning *Salvia divinorum* leaves and as a significant constituent of tobacco smoke^[1, 2]^. Many bioactive compounds contain 3HP as an important structural unit, and 3HP is widely used as a building block to synthesize drugs, herbicides, insecticides and antibiotics^[3-5]^. Large amounts of 3HP are synthesized each year, and there are many synthetic methods for 3HP, such as ruthenium-catalyzed ring-closing olefin metathesis^[6]^. The widespread use of 3HP has made its release to the environment inevitable, which may have serious implications for human health. While 3HP can be eliminated by physical and chemical methods, microbial biodegradation has been considered one of the most economical and effective approaches to remediating 3HP pollution.

Catabolism of pyridine, particularly the initial steps of the hydroxylation of monohydroxylated pyridines such as 2-hydroxypyridine (2HP), 3HP and 4-hydroxypyridine (4HP), has received much attention. In particular, numerous bacteria have been reported to use 2HP as the sole carbon and energy source to grow. Many intermediates have been identified, and metabolic pathways have been proposed for 2HP biodegradation. One pathway of 2HP biodegradation involves the formation of 2,5-dihydroxypyridine (2,5-DHP), which then proceeds through the maleamate pathway. The other pathway, involving the formation of 2,3,6-trixydroxypyridine first, produces a blue pigment (nicotine blue) in the medium. *Rhodococcus rhodochrous* PY11 was reported to use 2HP as the sole source of carbon and energy through the nicotine blue-production pathway. A gene cluster (*hpo*) has been characterized as being responsible for the catabolism of 2HP in strain PY11, and the initial hydroxylation of 2HP is catalyzed by a four-component dioxygenase (HpoBCDF)^[7]^. *Burkholderia* sp. MAK1 was also reported to degrade 2HP, but through the maleamate pathway. A gene cluster (*hpd*) was responsible for the degradation of 2HP in strain MAK1, and the 2-hydroxypyridine 5-monooxygenase is a soluble di-iron monooxygenase (SDIMO) encoded by a five-gene cluster *hpdABCED*^[8]^. Several strains have been reported to partially or completely degrade 3HP. *Achromobacter* sp. (G2 and 2L)^[9]^, *Pusillimonas* sp. 5HP^[10]^ and *Agrobacterium* sp. DW-1^[11]^ could use 3HP as a carbon and nitrogen source to grow. *Nocardia* Z1 was reported to slowly oxidize 3HP to pyridine-2,3-diol and pyridine-3,4-diol^[12]^, while *Achromobacter* 7N could convert 3HP to only 2,5-DHP and could hardly further metabolize the diol^[9]^.

Moreover, microbial degradation of 3HP, which has been proposed for several different strains, was thought to proceed via the maleamate pathway^[9, 12-14]^, 3HP →2,5-DHP → formate + maleamate → NH3 + maleate ↔ fumarate, and was recently confirmed in strain DW-1^[11]^. However, the genes and enzymes responsible for 3HP biodegradation have seldom been reported. Pyridine-2,5-diol dioxygenase has been partially purified and characterized in only strains G2 and 2L^[9]^. The pyridine-2,5-diol dioxygenase in these two strains required Fe^2+^ to restore full activity after purification, and the hydroxylases of strains G2 and 2L showed clear specificity because they produced only the *para*-substituted 2,5-DHP from 3HP. No enzymes responsible for the initial hydroxylation step of 3HP leading to the formation of 2,5-DHP have been reported to date.

In this study, the bacterial strain *Ensifer adhaerens* HP1 was isolated from soil and showed effective degradation and utilization of 3HP. We report the isolation and characterization of the 3HP catabolic pathway in *E. adhaerens* HP1. A gene cluster (*3hpd*) encoding the putative proteins required for 3HP biodegradation in this bacterium was discovered and characterized. The results of bioinformatics analysis, gene knockout and complementation of *hpdA*, and heterologous expression of HpdA suggest that multicomponent HpdA is involved in the transformation of 3HP to 2,5-DHP.

## Results and discussion

### Isolation of a bacterium capable of degrading 3HP

A bacterium was isolated from enrichments of soil with 3HP and was selected for detailed study. This strain could utilize 3HP as the sole source of carbon, nitrogen and energy to grow (Figure 1) and was designated HP1. The 16S rRNA sequence of strain HP1 exhibited the highest similarity (99%) to *E. adhaerens* strain Casida A (accession number: CP015882.1). Therefore, strain HP1 was identified as *E. adhaerens* HP1. During the growth of strain HP1 on 3HP-containing media, a green pigment accumulated and gradually turned dark brown. This phenomenon has been reported in strains *Pusillimonas* sp. 5HP^[10]^, *P. putida* S16^[15]^, and *Agrobacterium* sp. S33^[16]^. All these strains converted the corresponding substrate to 2,5-DHP, and this color change indicated the formation of 2,5-DHP. 2,5-DHP was detected in the supernatant of strain HP1 by LC-MS analysis (Figure S1), suggesting that the degradation of 3HP in strain HP1 occurs through the maleamate pathway (Figure 2B). Moreover, strain HP1 utilized only 3HP but not the other two hydroxypyridine isomers, 2HP and 4HP (data not shown).

**Figure 1.**
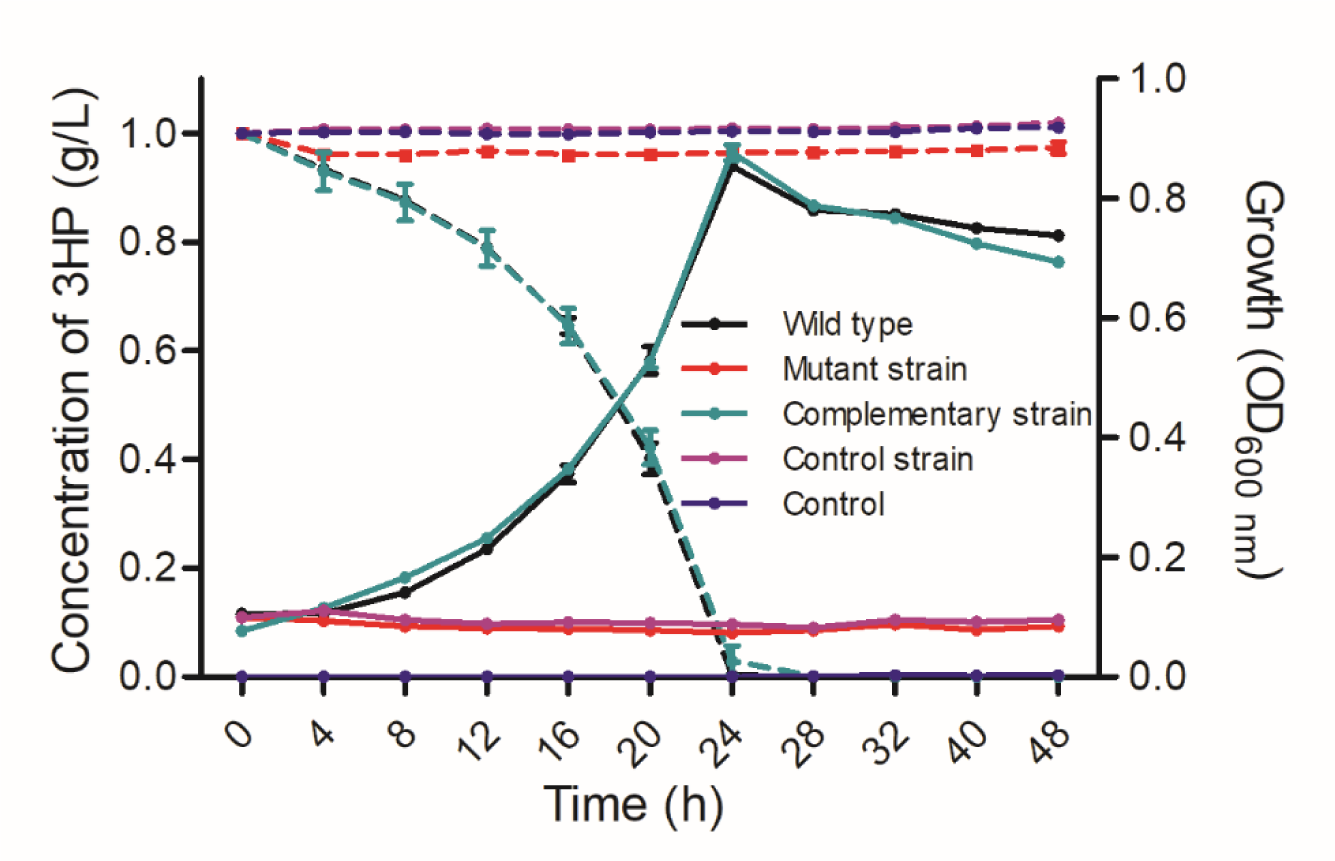
Cell growth of and 3HP degradation by strain HP1 and its plasmid transformants. The solid and broken lines represent cell growth and 3HP degradation, respectively. The results presented in these histograms are the means of three independent experiments, and the error bars indicate the standard error.

**Figure 2.**
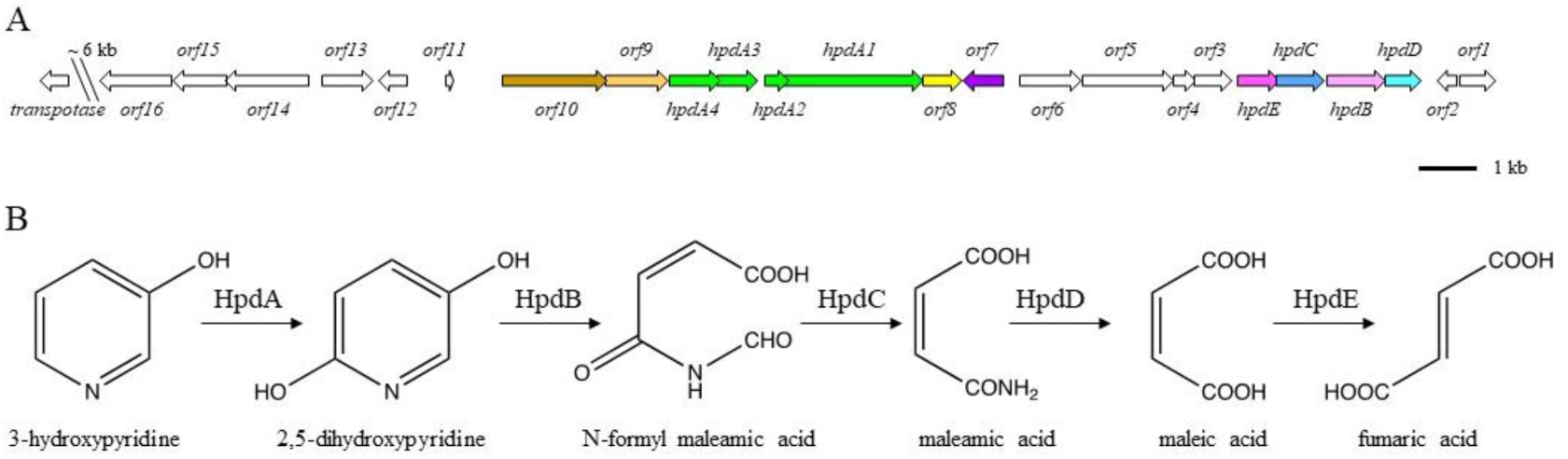
(A) Genetic organization of the putative 3HP degradation gene cluster *3hpd* in *E. adhaerens* HP1. The diagram shows the arrangement of genes that BLAST analysis suggests are involved in 3HP catabolism. The arrows indicate the size and direction of transcription of each gene. *hpdA* is a putative 3-hydroxypyridine dehydrogenase; *hpdBCDE* are putative 2,5-dihydroxypyridine dioxygenase, N-formyl maleamic acid deformylase, maleamate amidase, and maleate isomerase, respectively, which are responsible for the maleate pathway of 2,5-dihydroxypyridine metabolism. *Orf1-orf16* are genes flanking the putative 3HP metabolic genes. The predicted functions of these genes are summarized in Table 1. (B) Proposed catabolism of 3HP by *E. adhaerens* HP1.

**Table 1.**
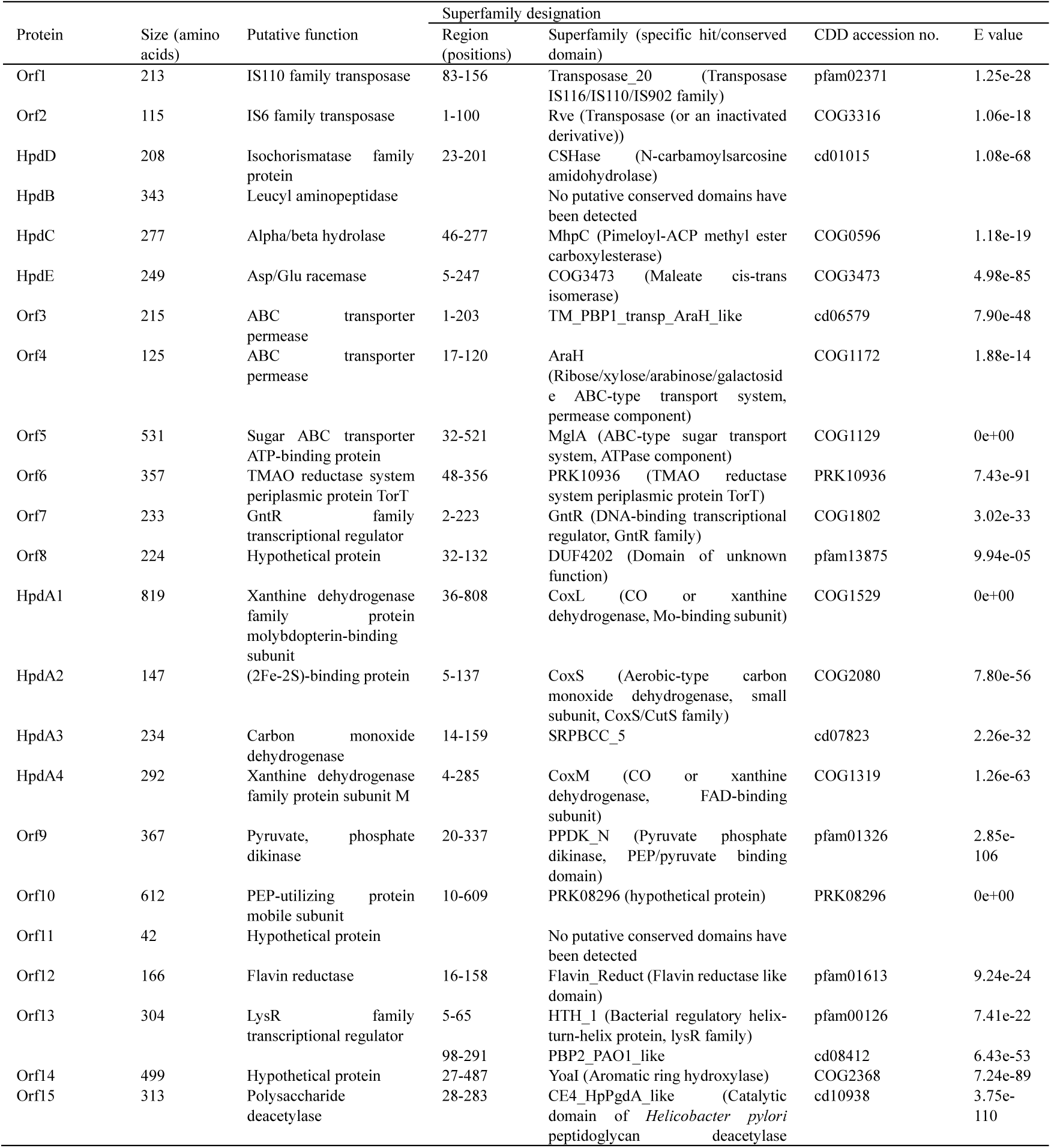
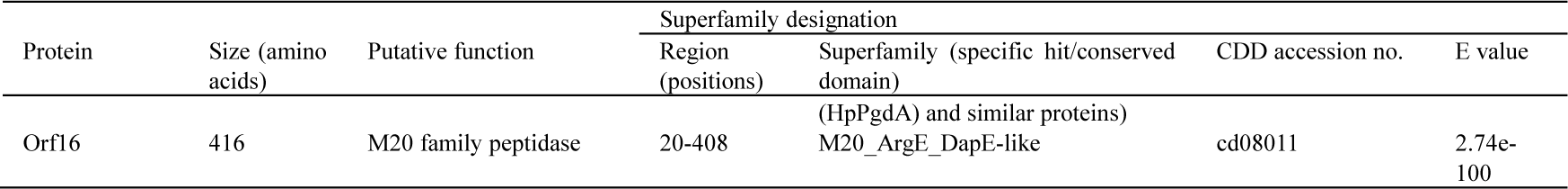
Functional annotations of hypothetical Hpd proteins

### A putative 3HP metabolism gene cluster is present in the genome of strain HP1

After performing a BLAST analysis against the genome of strain HP1 using previously known metabolic genes involved in the maleamate pathway, we found a putative metabolic gene cluster, *hpdDBCE*, in the genome of strain HP1 (Figure 2A). Analysis of the pyridine ring hydroxylase showed that there were four sets of putative genes that may be responsible for the initial metabolism of 3HP. Among them, a four-component gene, *hpdA1A2A3A4*, was contiguous with the putative maleamate pathway metabolic gene cluster. We predicted that the putative maleamate pathway metabolic gene and the four-component gene were the most likely components of the 3HP-degrading operon.

The cluster consists of 24 genes, *hpdA1*, *hpdA2*, *hpdA3*, *hpdA4*, *hpdB*, *hpdC*, *hpdD*, *hpdE*, and *orf1*-*orf16* (Figure 2A), and we designated this cluster *3hpd*. The functional annotations of hypothetical Hpd proteins are listed in Table 1. *orf1* and *orf2* encode transposases. The amino acid sequences of *orf3*-*orf5* showed similarity to the bacterial ABC transporter. Gene *orf7* in *3hpd* encodes a potential GntR family DNA-binding transcriptional regulator. Moreover, it showed that the transcription of *orf3*-*orf5* was induced by 3HP (Figure 3). This allowed us to presume that *orf3*-*orf5* may be involved in the transport of 3HP, but this requires further research. BLAST analysis based on *3hpd* showed that *hpdA* had few homologous genes, which showed relatively low nucleotide sequence similarities (<75%), except for one uncharacterized gene cluster in *Rhizobiales* bacterium isolate AFS066724 (sequence accession number: UCDB01000016.1), with nucleotide sequence similarity >95.92%. It was reported that bacterial species display a wide range of variation in their total G+C content, and the genes in a certain species’ genome share similar nucleotide sequence patterns^[17, 18]^. The G+C content of the whole genome of strain HP1 (7.18 Mb) was 62.07%, whereas the G+C content of the plasmid (334.2 kb) containing the *3hpd* gene cluster and the *3hpd* gene cluster alone (calculated for genes between *hpdD* and *hpdA4*, 13.2 kb) were 60.40% and 60.30%, respectively. We observed that the G+C contents of the plasmid and the *3hpd* gene cluster were obviously lower than that of the whole genome. In addition, considering the mobile element remnants flanking the *3hpd* gene cluster (Figure 2A), which is a direct indication of a foreign origin of the sequence flanked by mobile elements, the *3hpd* gene cluster was probably introduced through horizontal gene transfer (HGT) to the genome of strain HP1.

**Figure 3.**
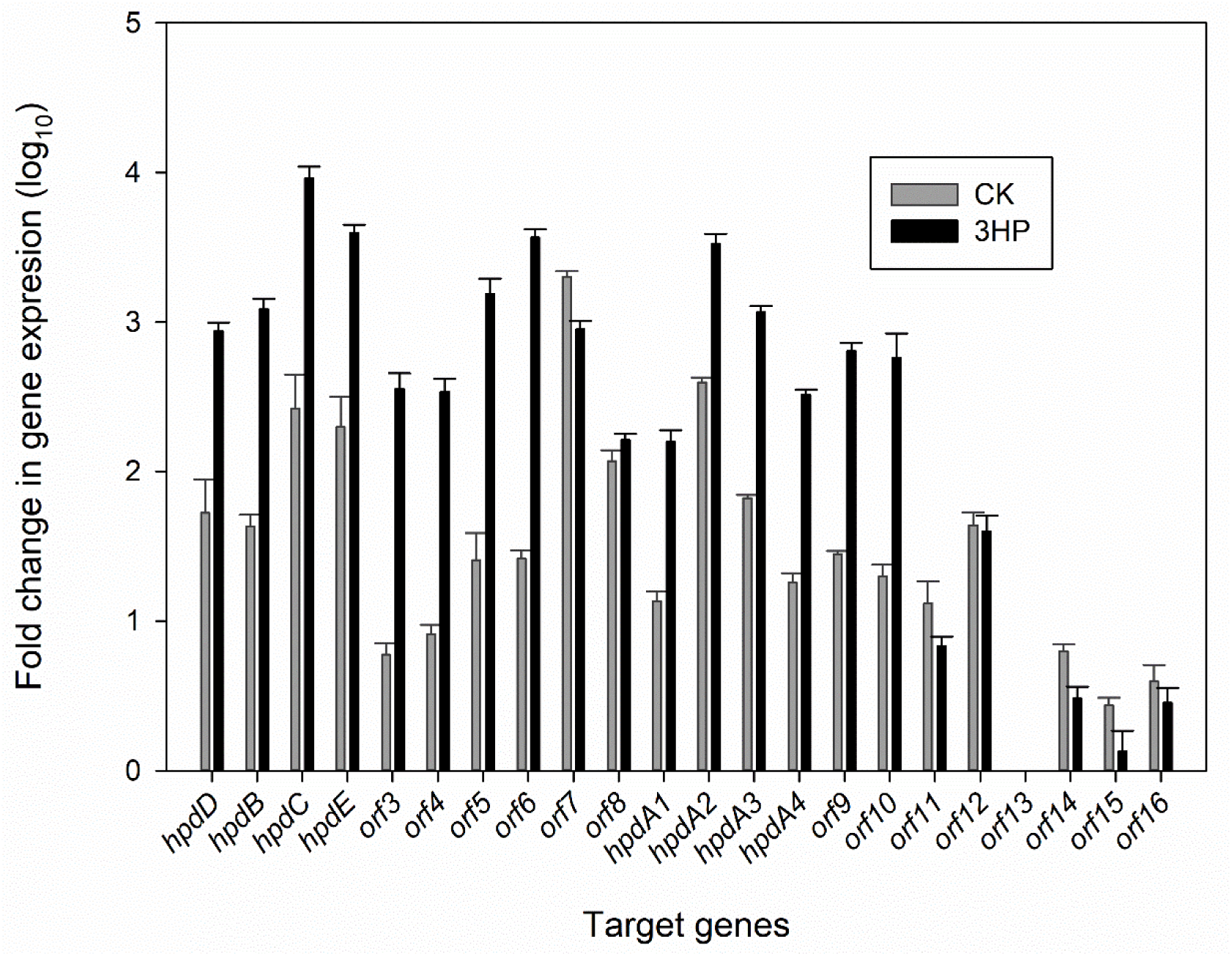
Transcriptional analysis of the *3hpd* gene cluster. RT-qPCR analysis of target gene transcripts produced in *E. adhaerens* HP1 grown with (black bars) or without (gray bars) 3HP. The expression levels of the *3hpd* genes were normalized to the 16S rRNA expression level and are expressed as the fold change in expression in cells. The results presented in these histograms are the means of four independent experiments, and error bars indicate the standard error.

### Transcription of putative 3HP-degrading genes in *3hpd* is induced by 3HP

To elucidate the correlation between 3HP degradation and putative 3HP-degrading genes in *3hpd* and other candidate 3HP dehydrogenase genes, the expression levels of putative target genes involved in the 3HP degradation of strain HP1 were estimated using reverse transcription quantitative PCR (RT-qPCR) and the 2^ΔΔCT^ method with or without 3HP supplementation in minimal salt medium (MSM). As shown in Figure 3, the expression levels of *hpdA1*, *hpdA2*, *hpdA3*, *hpdA4*, *hpdB*, *hpdC*, *hpdD*, and *hpdE* in *3hpd* in experimental group were more than 10-times higher than the levels of the corresponding genes in the control group, suggesting that the expression of these genes was induced by 3HP or other 3HP degradation intermediates, while other candidate 3HP dehydrogenase genes exhibited opposite results or even no transcription in the tested growth conditions (Figure S2).

### The *hpdA* genes encode the 3HP dehydrogenase

The first step in the metabolism of 3HP is the hydroxylation of C6 on the pyridine ring to generate 2,5-DHP. The hydroxylation of N-heterocyclic aromatic compounds consisting of a pyridine ring, such as nicotine to 6-hydroxynicotine and nicotinic acid to 6-hydroxynicotinic acid is usually catalyzed by multicomponent molybdopterin-containing dehydrogenases^[19, 20]^. A BLAST homology search of genes in *3hpd* against database sequences revealed that the genes *hpdA1*, *hpdA2*, *hpdA3*, and *hpdA4* shared the highest amino acid sequence identity (33%-52%) with the respective subunits of the three-component molybdopterin-containing dehydrogenases, such as nicotine dehydrogenase (NdhLSM)^[19]^ and quinoline 2-oxidoreductase (QorLSM)^[21]^. *hpdA1* was predicted to contain binding domains for the molybdopterin cytosine dinucleotide (MCD), *hpdA4* contained a flavin adenine dinucleotide (FAD) binding domain and specific FAD-binding site of members of the xanthine dehydrogenase/oxidase family (CODH), *hpdA2* contained two predicted [Fe-S] clusters, and *hpdA3* contained an SRPBCC ligand-binding domain (Figure 4). Therefore, *hpdA1*, *hpdA2*, *hpdA3*, and *hpdA4* were predicted to encode a four-component molybdopterin-containing dehydrogenase that catalyzes the initial step in the 3HP metabolic pathway.

**Figure 4.**
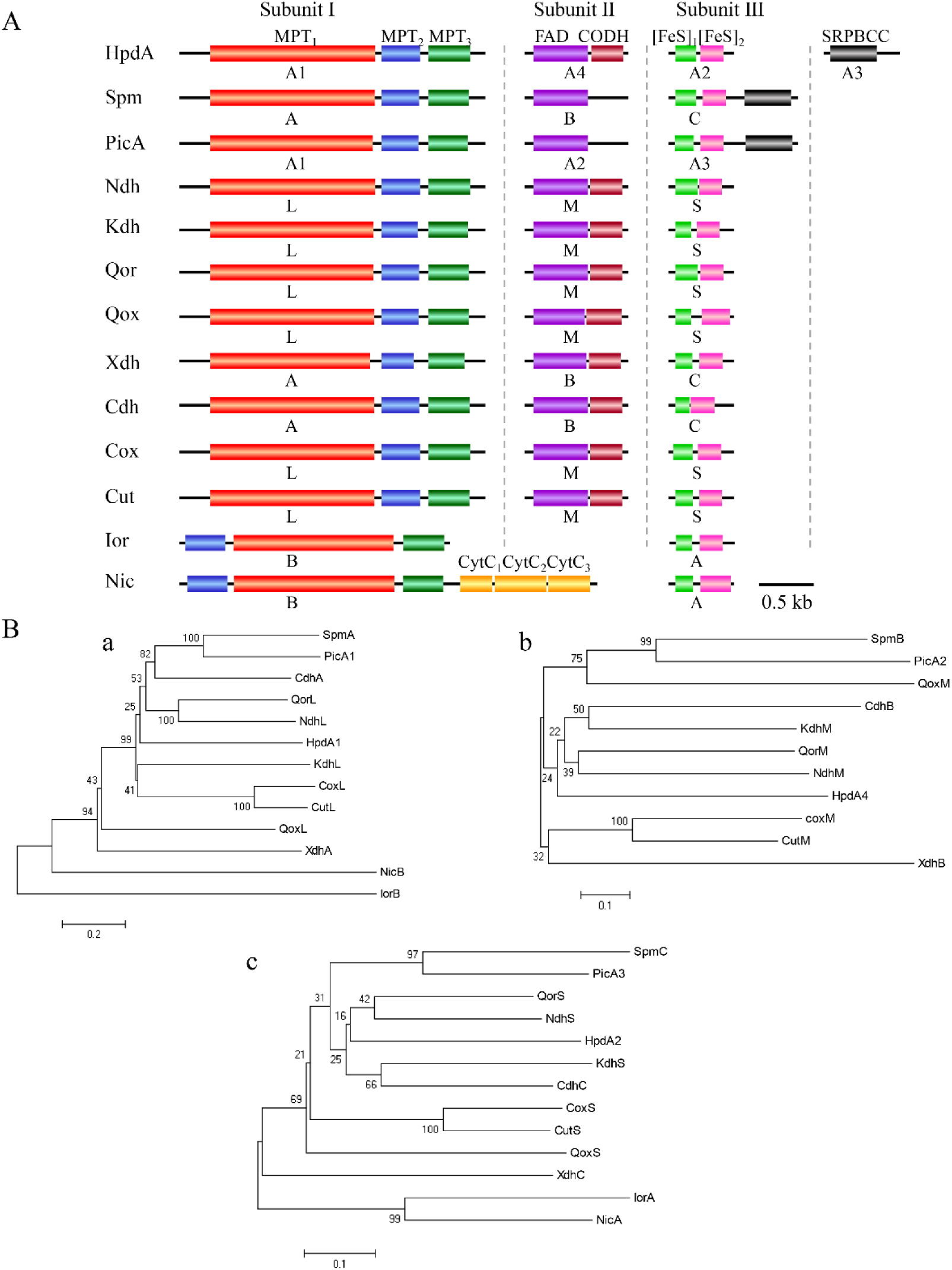
Bioinformatic analysis of HpdA. (**A**) Molecular architecture of several typical multicomponent molybdenum-containing hydroxylases. Subunits Ⅰ, Ⅱ, and Ⅲ are MCD-, FAD-, and two [Fe-S] cluster-containing components, respectively. HpdA, 3-hydroxypyridine dehydrogenase from *E. adhaerens* HP1; SpmABC (GenBank accession numbers AEJ14617 and AEJ14616), 3-succinoylpyridine dehydrogenase from *P. putida*; PicA (KY264362), picolinic acid dehydrogenase from *Alcaligenes faecalis* JQ135; NdhLSM (YP 007988778, YP 007988779, and YP 007988780), nicotine dehydrogenase from *Arthrobacter nicotinovorans*; KdhLMS (YP 007988766, YP 007988771, and YP 007988772), ketone dehydrogenase from *A. nicotinovorans*; QorLSM (X98131), quinoline 2-oxidoreductase from *P. putida* 86; QoxLMS (AJ537472), quinaldine 4-oxidase from *Arthrobacter ilicis* Ru61a; XdhABC (BAE76932.1, BAE76933.1, and BAE76934.1), xanthine dehydrogenase from *E. coli* W3110; CdhABC (ADH15879.1, ADH15880.1, and ADH15881.1), caffeine dehydrogenase *Pseudomonas* sp. strain CBB1; CoxLSM (X82447), carbon monoxide dehydrogenase from *Oligotropha carboxidovorans*; CutLSM (AAD00363.1, AAD00362.1, and AAD00361.1), carbon monoxide dehydrogenase from *Hydrogenophaga pseudoflava*; IorAB (CAA88753 and CAA88754), isoquinoline 1-oxidoreductase from *Brevundimonas diminuta* 7; NicAB (PP3947 and PP3948), nicotinate hydroxylase from *P. putida* KT2440. The letters below the proteins indicate the subunit names of the corresponding proteins. The conserved domains are MPT1, MPT2 and MPT3, which are the three motifs for binding to the MCD cofactor; FAD, consensus FAD-binding site; CODH, specific FAD-binding site of members of the xanthine dehydrogenase/oxidase family; [FeS]1, binding site for the ferredoxin-like [2Fe-2S] cluster; [FeS]2, binding site for the second [2Fe-2S] cluster; SRPBCC, SRPBCC ligand-binding domain; CytC1, CytC2 and CytC3, the three cytochrome c binding motifs. (**B**) Phylogenetic analysis of HpdA and related molybdenum-containing hydroxylases: HpdA1 and the Moco binding subunit of other enzymes (a); HpdA4 and the FAD-binding subunit of other enzymes (b); HpdA2 and the [2Fe-2S]-binding subunit of other enzymes (c). The phylogenetic trees were constructed using the neighbor-joining method (with 1,000 bootstraps) with MEGA software, version 6.06. The bar represents the number of amino acid substitutions per site.

To investigate the role of *hdpA* in the initial oxidation of 3HP, *hpdA1A2* was disrupted. Then, the ability of the mutant strain HP1Δ*hpdA* to metabolize 3HP was analyzed. The disruption of *hpdA1A2* genes abolished the growth of the mutant HP1Δ*hpdA* in 3HP-containing medium (Figure 1) but did not affect its growth in 2,5-DHP-containing medium (data not shown). After *trans* complementation of *hdpA1A2* to HP1Δ*hpdA*, the ability to utilize 3HP to grow was restored. The complementation of *hpdA1* alone could not recover the degradation ability (data not shown). To confirm the role of *hpdA* genes, they were cloned and expressed in plasmid pRK415-*hpdA*-*3.* When plasmid pRK415-*hpdA*-*3* was transferred to *E. adhaerens* ZM04, which is not able to utilize 3HP, the culture media turned green and then gradually dark brown, suggesting the formation of 2,5-DHP. Results showed that the recombinant strain ZM04-*hpdA-3* acquired the ability to efficiently convert 3HP into a stoichiometric amount of 2,5-DHP (Figure 5). UV spectral analysis showed that 3HP decreased over time (absorbance peaks at 244, 275, and 311 nm, pH 7.5), with the formation of a 2,5-DHP peak (absorbance peaks at 228 and 321 nm, pH 7.5) (Figure 6). LC-MS analysis also confirmed that the product was 2,5-DHP [molecular ion peak (M+H)^+^ 112.3680] (Figure S3). The recombinant ZM04-*hpdA* could convert 1 mM 3HP into equivalent amounts of 2,5-DHP (Figure 5), suggesting that 3HP was completely transformed into 2,5-DHP and that 2,5-DHP was the sole product of 3HP.

**Figure 5.**
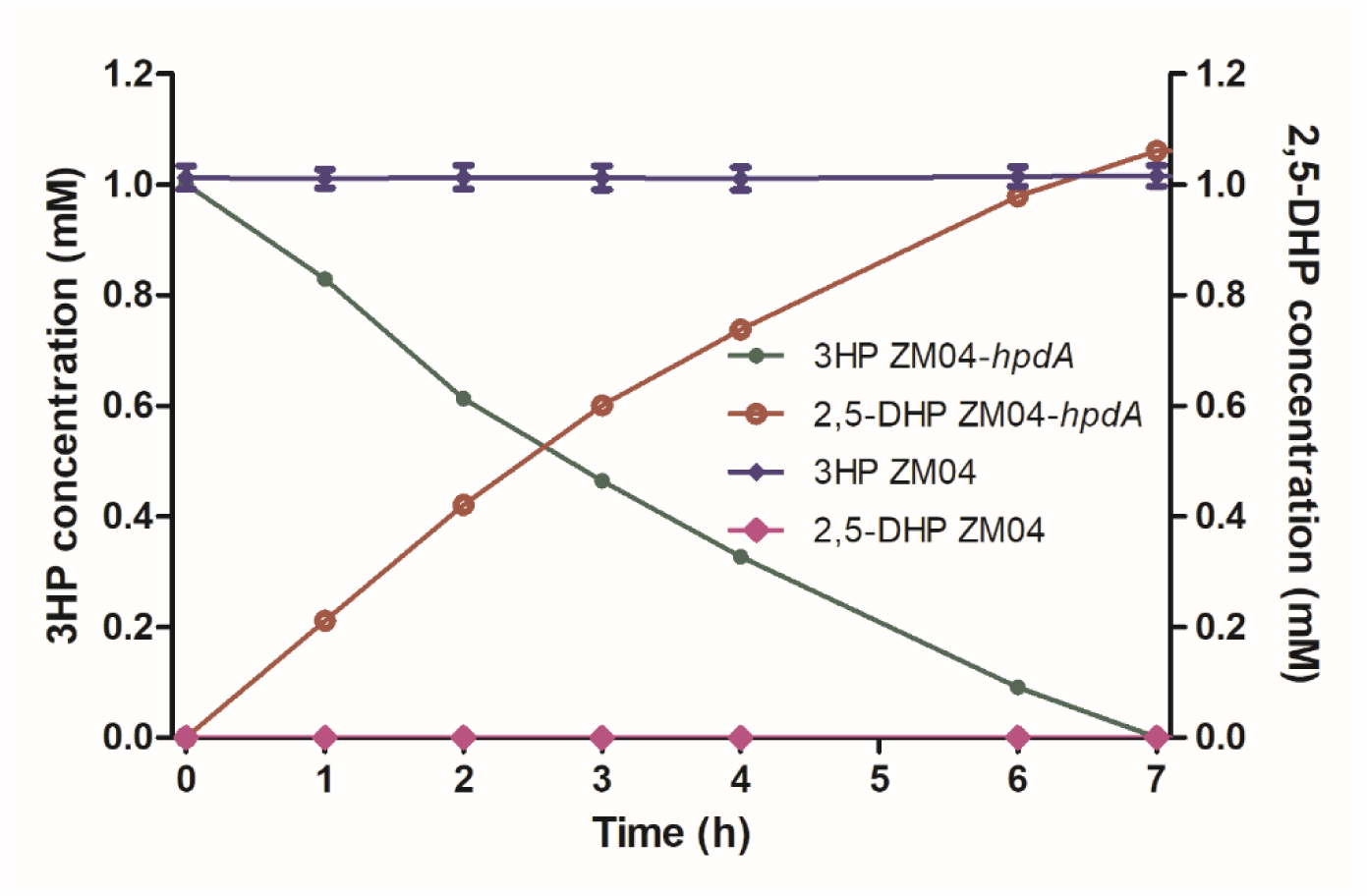
Time course of the conversion of 3HP to 2,5-DHP by recombinant strain ZM04-*hpdA*. The control group with strain ZM04 did not show transformation of 3HP, and no product was formed.

**Figure 6.**
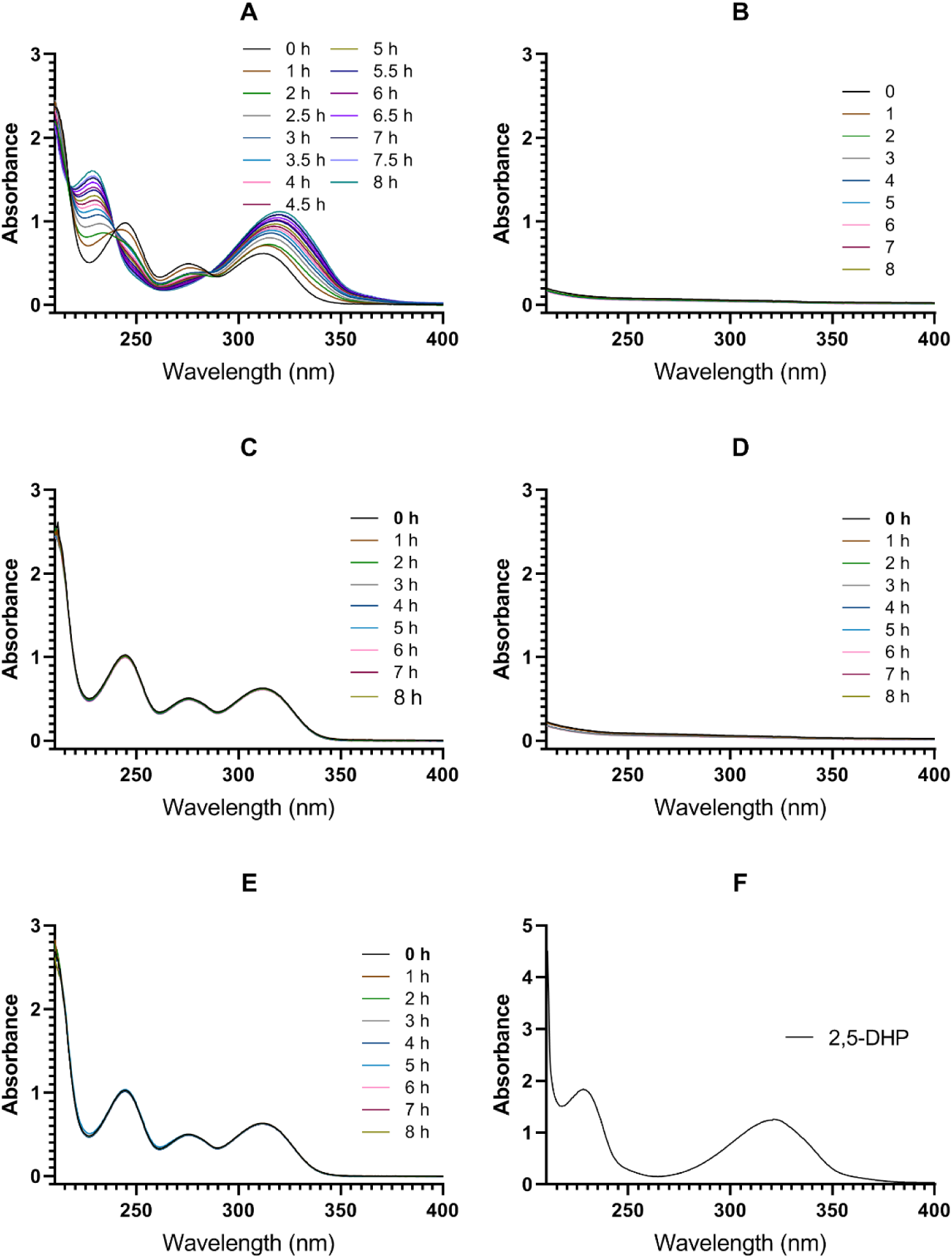
Detection of the catalytic activity of *hpdA* in strain ZM04. UV spectrum for the conversion of 3HP by transformant ZM04 pRK415-*hpdA-3* with substrate 3HP (A), ZM04 pRK415-*hpdA-3* without substrate 3HP (B), ZM04 pRK415 with substrate 3HP (C), ZM04 pRK415 without substrate 3HP (D), and substrate 3HP only (E) and UV spectrum of 0.025 mg/mL of 2,5-DHP standard (F). Samples in A-E were diluted 5 times before detection. Each group was analyzed three times independently, and the results of each group showed a similar trend. The best one is shown.

When truncated HpdA lacking *hpdA3* or *hpdA4* was constructed and expressed from plasmids pRK415-*hpdA-4* and pRK415-*hpdA-5,* respectively, these two truncated enzymes did not show 3HP dehydrogenase activity, and no 2,5-DHP or color change in the culture media was detected during incubation (data not shown). These results showed that all of the components encoded by *hpdA1*, *hpdA2*, *hpdA3* and *hpdA4* are essential for the function of HpdA. In summary, our data demonstrate that *hpdA* encodes the dehydrogenase that catalyzes the oxidation of 3HP.

### Prediction and analysis of transmembrane domain and signal peptide in *hpdA3*

HpdA3 contained a SRPBCC ligand-binding domain. It was reported that SRPBCC domains have a deep hydrophobic ligand-binding pocket that can bind diverse ligands and spans all three kingdoms of life, but its function remains unclear. Moreover, SRPBCC-containing genes play a wide range of roles, including strawberry fruit ripening^[22]^, regulation of anthocyanin accumulation^[23]^, encoding type II secretion chaperones^[24]^, functioning in the stabilization of mRNA^[25]^, catalyzing the free radical reactions of polyunsaturated fatty acids^[26]^, and α-pyridone ring formation^[27]^. Hydroxylases that hydroxylate the C6 of pyridine, such as 3-succinoylpyridine dehydrogenase (SpmABC)^[28]^ and picolinic acid dehydrogenase (PicA)^[29]^, include SRPBCC-containing components and are fused with [2Fe-2S] clusters, whereas most pyridine hydroxylases, such as NdhLSM and KdhLMS, have no SRPBCC domain (Figure 4A). In the case presented here, the SRPBCC domain (encoded by *hpdA3*) was not fused to the HpdA2 protein (containing the [2Fe-2S] cluster) but appeared in a separate protein, with an adjacent [2Fe-2S]-binding protein and FAD-binding subunit (Figures 2A and 4A). In addition, HpdA1, HpdA2 and HpdA4 showed relatively low evolutionary relationships with other molybdenum-containing hydroxylases (Figure 4B). The origin of this interesting and unique feature observed in HpdA is mysterious, and the function of this SRPBCC domain in HpdA deserves further study.

Analysis of the amino acid sequence of the *hpdA3* gene, encoding a carbon monoxide dehydrogenase, from strain HP1 using TMHMM, HMMTOP, TMpred and SignalP-5.0 predicted the presence of a TAT signal peptide (Sp1) (amino acids [aa] 138-176; likelihood, 0.913), followed by a transmembrane domain (Tmd) (aa 216-233). In addition, a different Sec signal peptide (Sp2) was also predicted by SignalP-5.0 (aa 155-176; likelihood, 0.8819). The predicted SRPBCC conserved domain of HpdA3 was located at aa 14-159. The 22 amino acid residues at the C-terminus of the SRPBCC domain were included in the Sp1 sequence. A previous study showed that some of the bacterial hydroxylases, such as nicotine dehydrogenase NDH in *A. nicotinovorans* pAO1^[19]^, CO dehydrogenase in *P. carboxydovorans* strain OM5^[30]^, and carbon monoxide dehydrogenase (CODH) in the eubacterium *Oligotropha carboxidovorans*^[31]^, were membrane associated. Bioinformatics analysis of the full-length HpdA3 predicted the presence of a signal peptide and transmembrane segment, which led us to presume that the protein encoded by *hpdA3* may function in the localization of HpdA. To confirm the roles of HpdA3 in the localization of HpdA, the full-length HpdA3 was fused to the N terminus of green fluorescent protein (GFP) and expressed in *Escherichia coli*. Confocal microscopy analysis of *E. coli* cells expressing GFP showed that the green fluorescent signal was uniformly dispersed in the cytoplasm, whereas the cells expressing HpdA3-GFP showed distinct uneven green fluorescence throughout the cell (Figure 7). Although the results for infusion proteins did not show typical membrane-associated localization, HpdA3 had a significant impact on the localization of infusion proteins. Based on these observations, we speculate that HpdA3 is probably essential for the localization of HpdA. SRPBCC-containing PicA3 and SpmC did not contain a signal peptide or transmembrane sequence, whereas HpdA3 contained both a signal peptide and Tmd domain. Therefore, in addition to the separate SRPBCC domain in HpdA, the signal peptide and the transmembrane domain in HpdA3 also distinguish HpdA from SpmABC and PicA.

**Figure 7.**
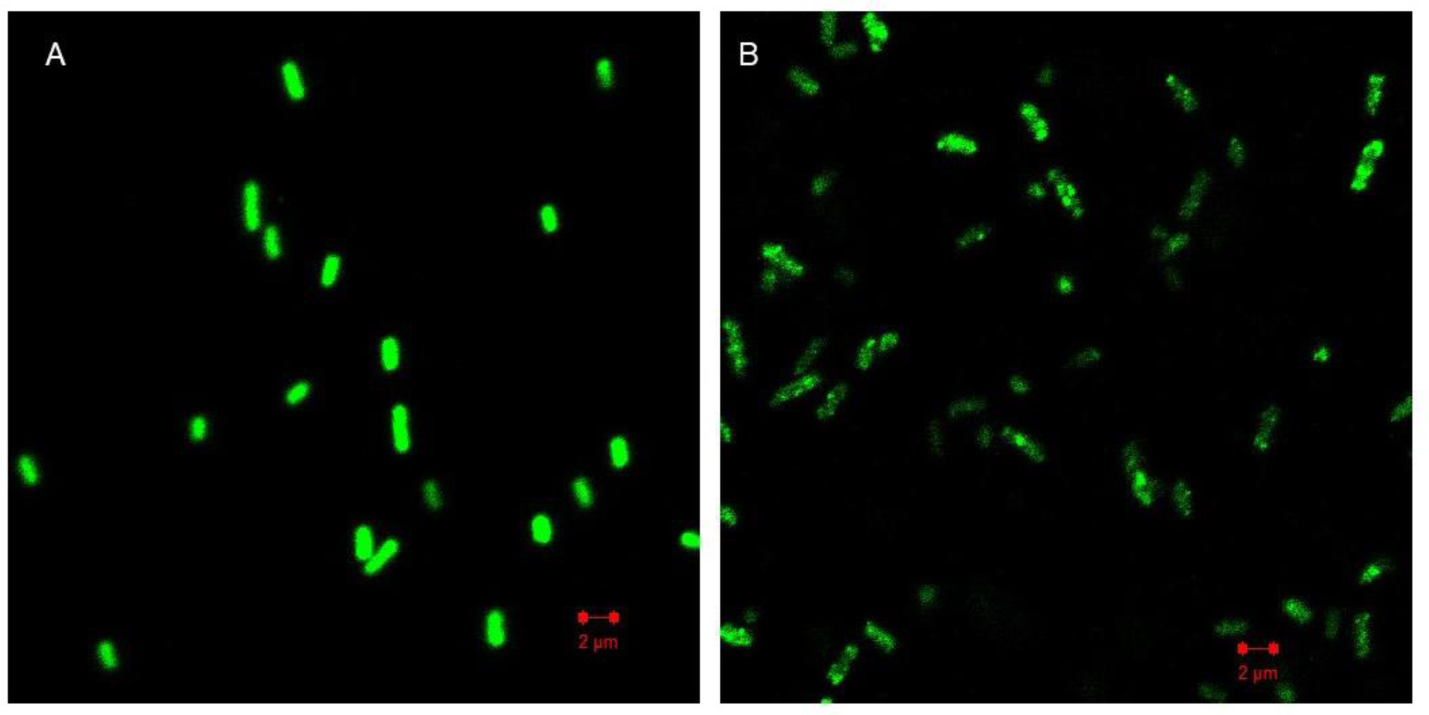
Localization of GFP by confocal microscopy. *E. coli* BL21 (DE3)-pGFPe (A) and *E. coli* BL21 (DE3)-pGFPe*hpdA3* (B).

### A phosphoenolpyruvate (PEP)-utilizing protein mobile subunit and pyruvate-phosphate dikinase are essential for 3HP conversion

To confirm the role of HpdA genes, we constructed several heterologous expression plasmids, including pRK415-*hpdA-1*, pRK415-*hpdA-2*, pRK415-*hpdA-3*, and pRK415-*hpdA-6* (Table 2). Plasmid pRK415-*hpdA-6* contains the *hpdA1A2A3A4* cluster, but when this plasmid was transferred to strain ZM04, the recombinant strain ZM04-*hpdA-6* did not acquire the ability to convert 3HP. To determine whether the genes contiguous with *hpdA* are also required for the function of HpdA, we constructed plasmid pRK415-*hpdA-1*, which contains *hpdA1A2A3A4*, as well as *orf7* and *orf8* at the 3-terminus and *orf9* and *orf10* at the 5-terminus of *hpdA*. As expected, when the plasmid was transferred to strain ZM04, the recombinant strain ZM04-*hpdA-1* obtained the ability to convert 3HP. To further confirm whether *orf7*-*orf8* or *orf9*-*orf10* are essential for HpdA function, we constructed plasmids pRK415-*hpdA-2* and pRK415-*hpdA-3*, with the former containing *hpdA1A2A3A4* and *orf7*-*orf8* and the latter containing *hpdA1A2A3A4* and *orf9*-*orf10*. Surprisingly, the latter imparted the recombinant strain ZM04-*hpdA*-*3* with the ability to transform 3HP as aforementioned, but the former did not. As described in Table 1, *orf10* encodes the PEP-utilizing protein mobile subunit, and *orf9* encodes pyruvate-phosphate dikinase. To further confirm the function of *orf9* and *orf10* in vivo, *orf9* and *orf10* were disrupted. The two mutants lost the ability to grow with 3HP as expected (data not shown), which showed that *orf9* and *orf10* are essential for HpdA function. What role do they play in the conversion of 3HP? In a previous study, CreH was found to encode a PEP-utilizing enzyme, and CreI encodes pyruvate-phosphate dikinase. The complex CreHI is a novel 4-methylbenzyl phosphate synthase that is responsible for the phosphorylation of the hydroxyl group of 4-cresol^[32]^. Another similar phosphorylation reaction is catalyzed by the phenylphosphate synthase reported in anaerobic phenol metabolism in *Thauera aromatica*, and it reported that the molecular and catalytic features of phenylphosphate synthase resemble those of phosphoenolpyruvate synthase^[33]^. According to these two previous studies, we speculated that *orf9* and *orf10* may be responsible for the conversion of 3HP to an intermediate compound, but during the experiment, we did not detect any product other than 2,5-DHP. The functions of *orf9* and *orf10* remain to be clarified.

**Table 2.**
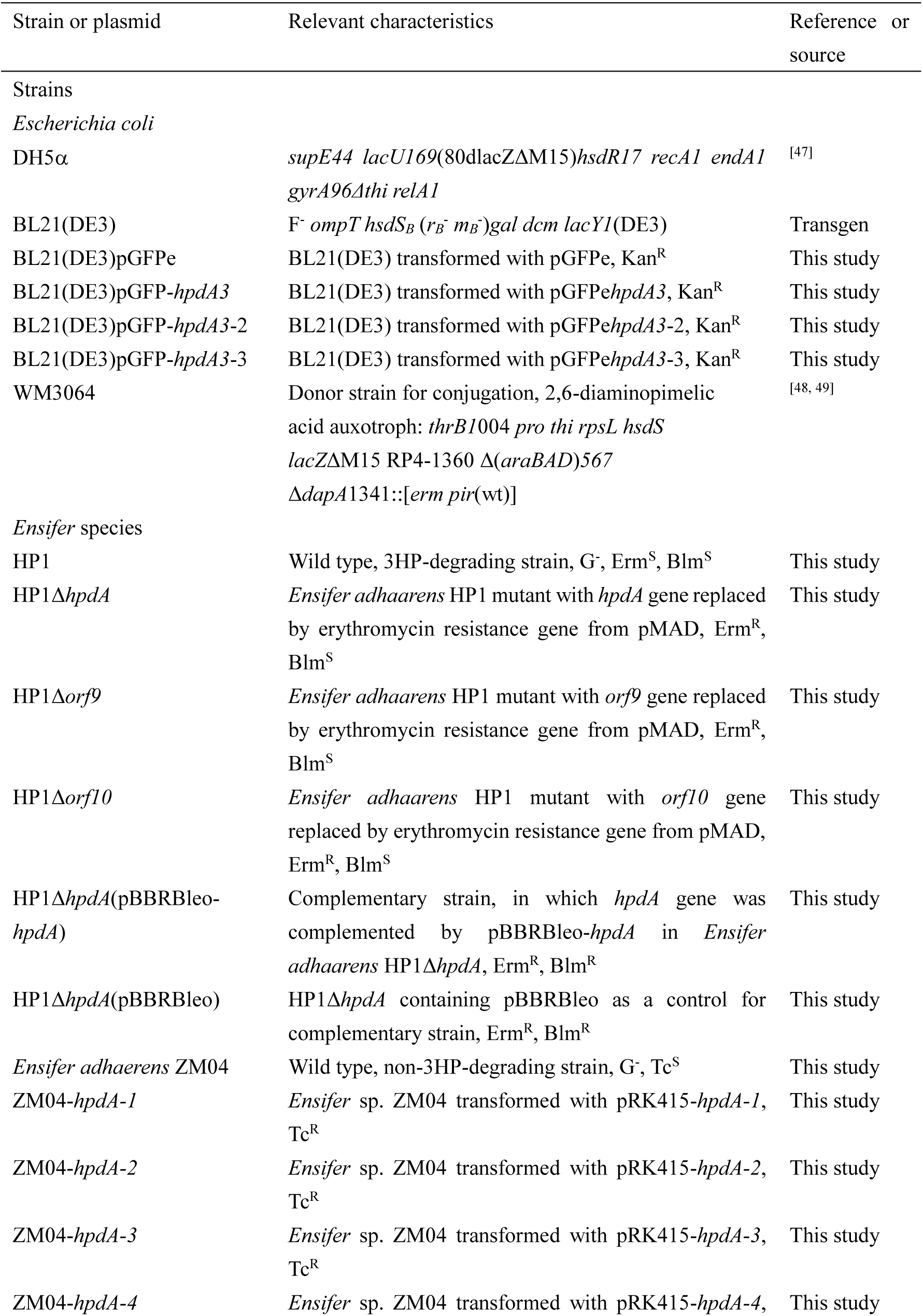
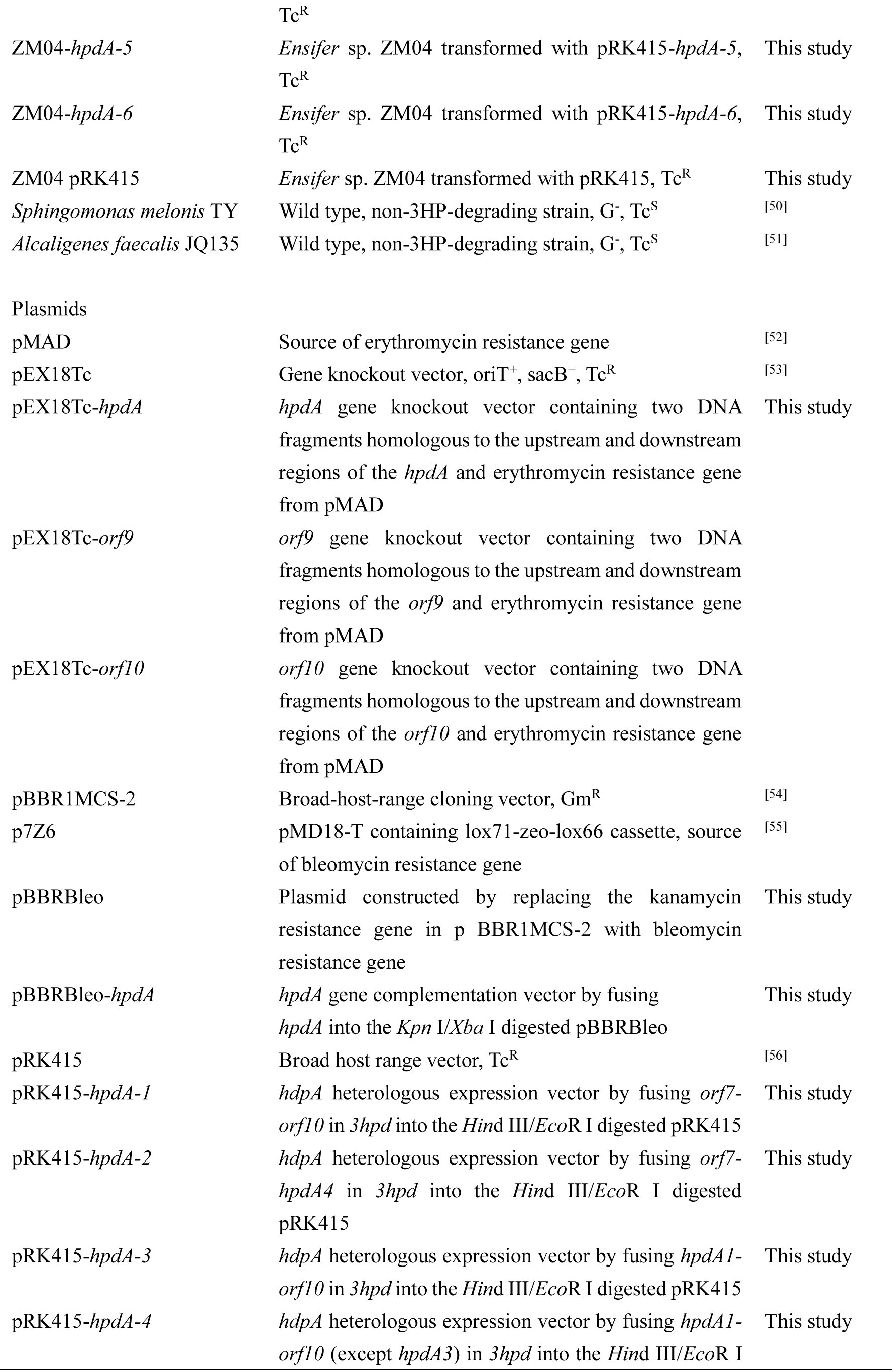
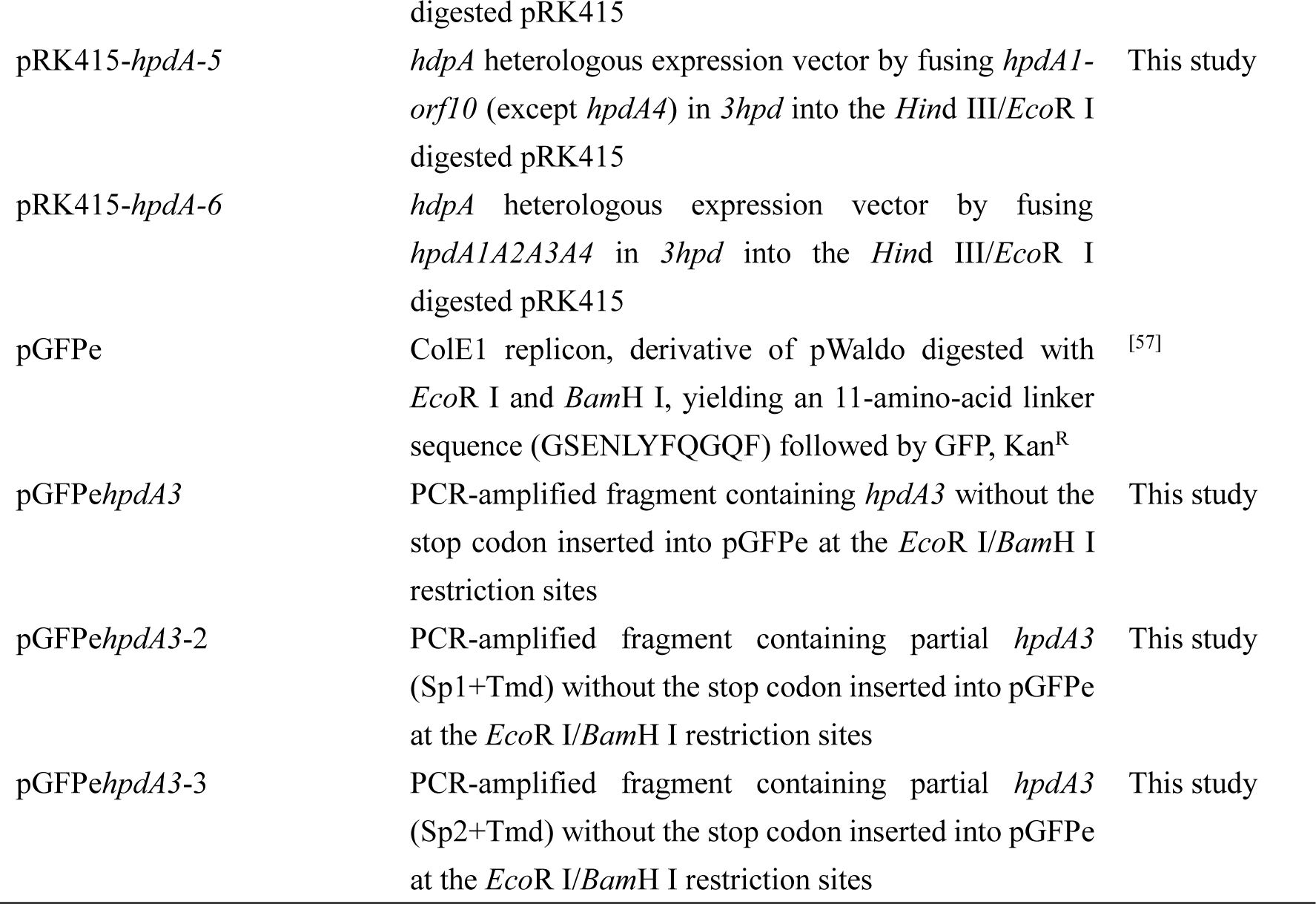
Strains and plasmids used in this study

### Distribution and diversity of *3hpd* genes in other bacteria

Eleven genome sequences of *E. adhaerens* strains (Casida A, OV14, RP12G, L18, ST2, WJB133 25_10, M78, AG1206, YX1, SD006, and X097) were available in NCBI, among which the genome sequence of strain Casida A was complete. The *3hpd* gene cluster was not identified in these genomes or other *Ensifer* species. Orthologous *3hpd* gene clusters were found in *Actinobacteria*, *Rubrobacteria*, *Thermoleophilia*, and *Alpha-*, *Beta-*, and *Gammaproteobacteria* (21 genera and 38 strains) (Tables S1 and S2). Most of the strains belong to the order *Burkholderiales* of the class *Betaproteobacteria* and the orders *Corynebacteriales*, *Geodermatophilales*, and *Pseudonocardiales* of the class *Actinobacteria*, including the following genera: *Pusillimonas*, *Acidovorax*, *Comamonas*, *Variovorax*, *Mycobacterium*, *Mycolicibacterium*, *Blastococcus*, *Geodermatophilus*, *Amycolatopsis*, and *Pseudonocardia*. Interestingly, some of the strains are degraders (e.g., *Acidovorax* sp. KKS102 is a polychlorinated-biphenyl-degrading strain^[34]^, *Pusillimonas noertemannii* BS8 can cooperate with another strain to degrade poly-γ-d-glutamic acid^[35]^, and *Mycobacterium* sp. MS1601 can oxidize branched polyols^[36]^), whereas some of the strains are unique (e.g., *Geodermatophilus sabuli* strain DSM 46844 is a γ-radiation-resistant actinobacterium^[37]^, *Amycolatopsis acidiphila* strain JCM 30562 is a member of the genus *Amycolatopsis* inhabiting acidic environments^[38]^, and *Salinicola peritrichatus* strain JCM 18795 was isolated from deep-sea sediment^[39]^).

The genetic organization of the *3hpd* gene clusters in these bacteria was highly diverse (Figure 8). The *hpdA1A2A3A4* genes were generally contiguous, except for orthologous genes in the class Actinobacteria and two orthologous genes in the class Alphaproteobacteria. In most genera, *orf9* and *orf10* were also contiguous and adjoined *hpdA4*, except in *Gaiella occulta* strain F2-233 and *Solirubrobacterales* bacterium 70-9 SCNpilot. Interestingly, the *3hpd* genes in the class Alphaproteobacteria, class Gammaproteobacteria, class Actinobacteria and family Alcaligenaceae (belonging to class Betaproteobacteria) were diverse, while the *3hpd* genes in the family Comamonadaceae (belonging to the class Betaproteobacteria) were highly conserved.

**Figure 8.**
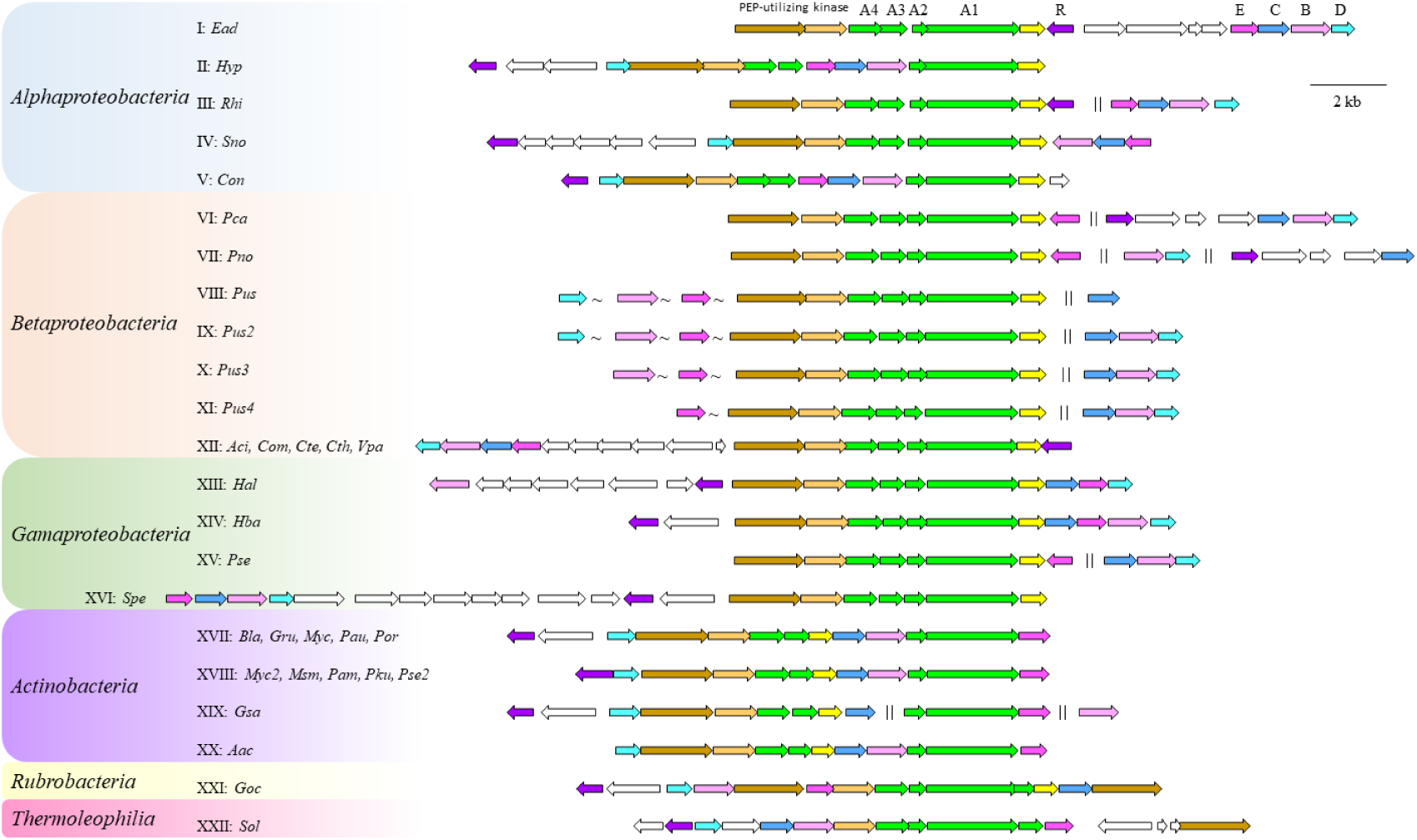
Predicted 3HP catabolism gene clusters in bacterial genomes. Ⅰ to XXII: the 22 different 3HP catabolism loci. Abbreviations and representative strains are as follows: *Ead*, *Ensifer adhaerens* HP1; *Hyp*, *Hyphomicrobium* sp. 99; *Rhi*, *Rhizobiales* bacterium isolate AFS066724; *Sno*, *Starkeya novella* isolate S2; *Con*, *Confluentimicrobium* sp. EMB200-NS6; *Pca*, *Pusillimonas caeni* strain KCTC 42353; *Pno*, *Pusillimonas noertemannii* BS8; *Pus*, *Pusillimonas* sp. 17-4A; *Pus2*, *Pusillimonas* sp. L52-1-41; *Pus3*, *Pusillimonas* sp. isolate EAC49; *Pus4*, *Pusillimonas* sp. isolate SAT20; *Aci*, *Acidovorax* sp. KKS102; *Com*, *Comamonas* sp. A23 and *Comamonas* sp. Z1; *Cte*, *Comamonas testosteroni* I2 and *Comamonas testosteroni* NBRC 100989; *Cth*, *Comamonas thiooxydans* strain S44; *Vpa*, *Variovorax paradoxus* isolate S2; *Hal*, *Halomonas* sp. MES3-P3E; *Hba*, *Halomonadaceae* bacterium R4HLG17; *Pse*, *Pseudoxanthomonas* sp. SGD-5-1; *Spe*, *Salinicola peritrichatus* strain JCM 18795; *Bla*, *Blastococcus* sp. DSM 46838; *Gru*, *Geodermatophilus ruber* strain DSM 45317; *Myc*, *Mycobacterium* sp. MS1601; *Pau*, *Pseudonocardia autotrophica* strain NRRL B-16064; *Por*, *Pseudonocardia oroxyli* strain CGMCC 4.3143; *Myc2*, *Mycobacterium* sp. GA-2829; *Msm*, *Mycolicibacterium smegmatis* MKD8; *Pam*, *Pseudonocardia ammonioxydans* strain CGMCC 4.1877; *Pku*, *Pseudonocardia kunmingensis* strain DSM 45301; *Pse2*, *Pseudonocardia* sp. MH-G8; *Gsa*, *Geodermatophilus sabuli* strain DSM 46844; *Aac*, *Amycolatopsis acidiphila* strain JCM 30562; *Goc*, *Gaiella occulta* strain F2-233; *Sol*, *Solirubrobacterales* bacterium 70-9 SCN. The detailed genomic accession numbers and the gene locus tags are listed in Table S1 in the supplemental material. Identities (percent) and similarity (percent) of amino acid sequences between *3hpd* proteins of strain *E. adhaerens* HP1 and representative homologs are listed in Table S2.

## Conclusions

3HP, generated through tobacco burning or industrially synthesized, is released into the environment and is potentially toxic or otherwise unacceptable. The degradation and utilization of 3HP by microbes have been studied for decades. This study isolated a 3HP-degrading bacterium, *E. adhaerens* HP1, that can utilize 3HP as the sole source of carbon and nitrogen to grow via the maleamate pathway. We also revealed that the *3hpd* gene cluster may be responsible for the degradation of 3HP in strain HP1 and that a four-component dehydrogenase, *hpdA*, catalyzes the initial hydroxylation of 3HP to 2,5-DHP. The genetic pathway of 3HP to 2,5-DHP is reported here for the first time. In addition, the separate SRPBCC component with predicted Sp and Tmd domains suggested the unusual evolution of HpdA, which needs further study. Moreover, the participation of a PEP-utilizing protein and pyruvate-phosphate dikinase in the conversion of 3HP by HpdA makes the catalytic mechanism of HpdA unique but requires further research. In summary, this work shows that the four-component HpdA protein constitutes a previously uncharacterized and unique dehydrogenase.

## Materials and methods

### Chemicals and reagents

3HP (>98%) and 2,5-DHP were obtained from Aladdin (Shanghai, China). TransStart*®* FastPfu DNA Polymerase and 2×T5 Super PCR mix for fragment amplification were purchased from TransGen Biotech (Beijing, China) and Beijing Tsingke Biotech Co., Ltd. (Beijing, China). Restriction enzymes used for plasmid construction were purchased from Takara Biotechnology Co., Ltd. (Dalian, China). Antibiotics, 2,6-diaminopimelic acid (2,6-DAP), β-D-1-thiogalactopyranoside (IPTG), and other reagents were purchased from Shanghai Sangon Biotech Co., Ltd. (Shanghai, China). All reagents and solvents were of analytical or chromatographic grade. A plasmid extraction kit, gel extraction kit and DNA purification kit were obtained from Omega Bio-tek, Inc. (Norcross, GA, USA). Bacterial genomic DNA was extracted using the TIANamp Bacterial DNA Kit from Tiangen Biotech Co., Ltd. (Beijing, China).

### Bacterial strains, plasmids, media, and growth conditions

Microorganisms with the ability to utilize pyridine as the sole source of carbon, nitrogen and energy were obtained by selective culture in shaken flasks, as described by C. Houghton and R. B. Cain^[12]^. Soil (1 g) was added to 250 mL Erlenmeyer flasks containing 100 mL of MSM containing the following (per liter): K_2_HPO_4_, 1 g; KCl, 0.25 g; MgSO_4_•7H_2_O, 0.25 g (sterilized separately and added aseptically to the cooled medium before use); and trace-element solution, 1 mL. The pH was adjusted to 7.5 using HCl (1 M). The trace-element solution contained the following (per liter): FeSO_4_•7H_2_O, 40 mg; MnSO_4_•H_2_O, 40 mg; ZnSO_4_•7H_2_O, 20 mg; CuSO_4_•5H_2_O, 5 mg; CoCl_2_•6H_2_O, 4 mg; Na_2_MoO_4_•2H_2_O, 5 mg; CaCl_2_•2H_2_O, 0.5 mg; and NaCl, 1 g. The 16S rRNA gene of the isolated 3HP-degrading strain was amplified with the universal primer set 27F and 1492R. Strain HP1 (collection number, CGMCC 1.13748) and its plasmid transformants were grown aerobically in MSM supplemented aseptically with 0.1% (wt/vol) 3HP in 250 mL Erlenmeyer flasks at 30 ℃ and 200 rpm. *E. coli* strains DH5α and WM3064 used in this study were grown in LB medium at 37 ℃ and 200 rpm. When necessary, erythromycin, bleomycin and tetracycline were used at final concentrations of 100, 25 and 10 μg/mL, respectively. 2,6-DAP was used at a final concentration of 0.3 mM for *E. coli* WM3064 and its transformants. The strains and plasmids used in this study are summarized in Table 2, and the primers are listed in Table 3.

**Table 3.**
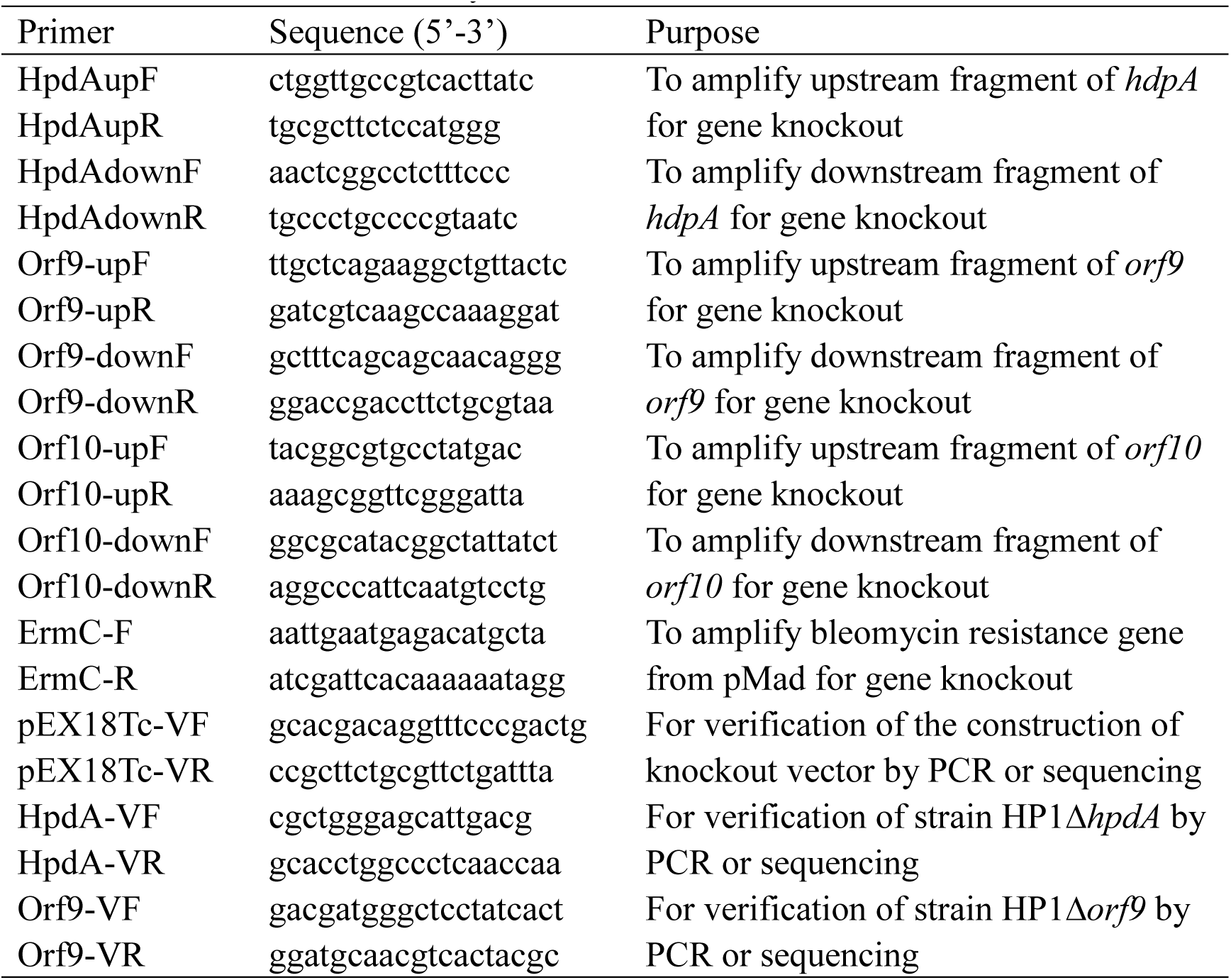
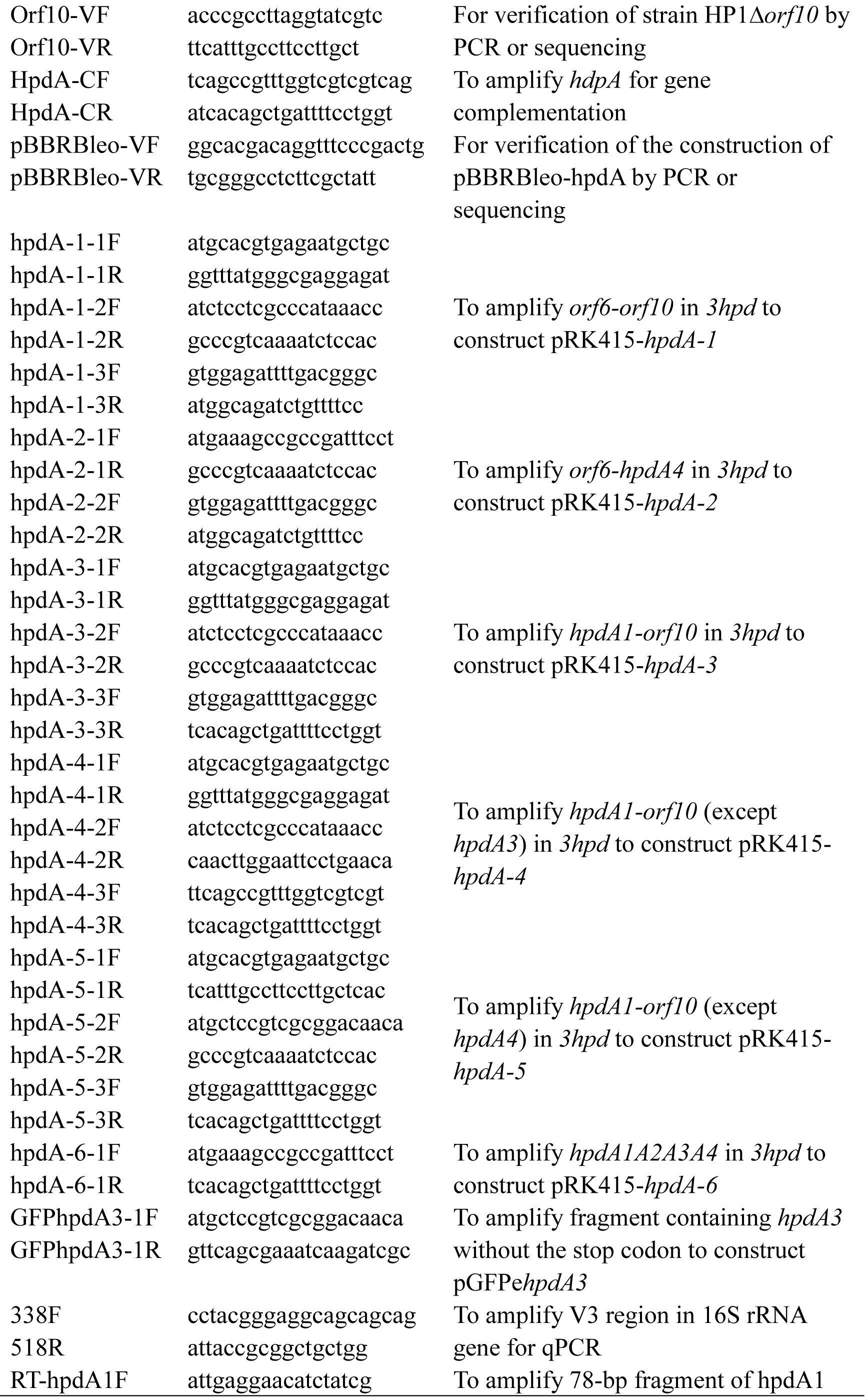
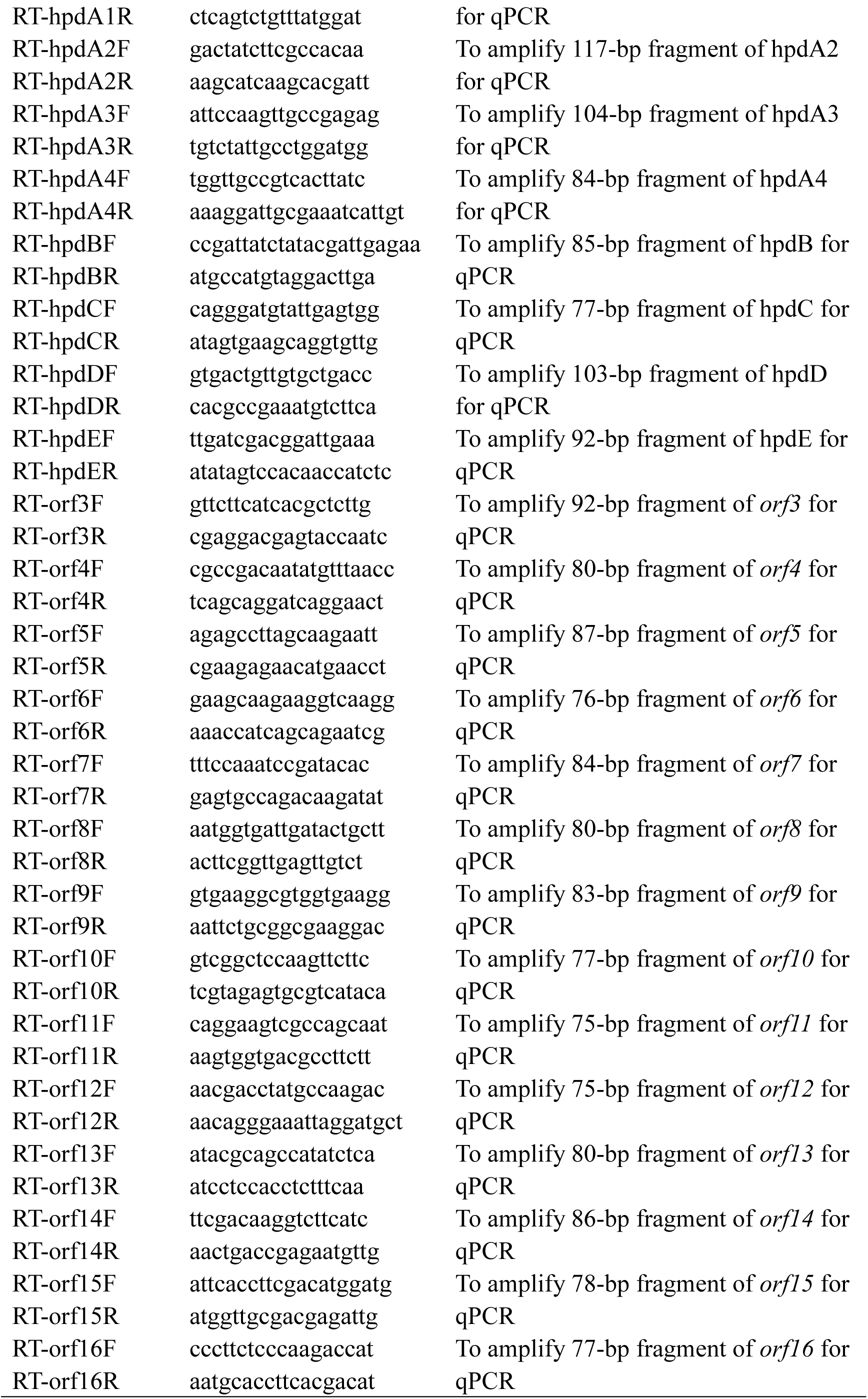
Primers used in this study

### Genome sequence analysis and prediction of 3HP metabolism-related genes in strain HP1

To identify putative genes involved in 3HP degradation in strain HP1, we conducted a BLAST analysis against the genome sequence of strain HP1 using known metabolic genes involved in the maleamate pathway, such as *ndpHFEG* in *Sphingomonas melonis* TY^[40]^ and *vppHFEG* in *Ochrobactrum rhizosphaerae* SJY1^[41]^. We also used some of the hydroxylases acting on the pyridine ring, such as *hpdABCED*^[8]^, *ndhLSM*^[19]^, *kdhLMS*^[42]^, *spmABC*^[28]^, and *nicAB*^[20]^, to predict the unknown dehydrogenase in initial metabolism of 3HP.

### RT-qPCR analysis

RT-qPCR analyses of strain HP1 cultures grown under different conditions for select genes were performed using specific primers (Table 3). For the experimental group, 3HP was used as the sole carbon and nitrogen source in MSM, while for the control group, glucose and ammonium sulfate were used as the carbon and nitrogen source in MSM, respectively. Total RNA was extracted from mid-exponential cultures of strain HP1 (both the control and experimental groups) using the RNAprep Pure Bacteria Kit (Tiangen Biotech, Beijing, China) according to the manufacturer’s protocol. All RNA samples were stored at -80 °C until further processing. To obtain the cDNA and remove potential traces of genomic DNA, RNA was reverse-transcribed into cDNA using random hexamer primers and the PrimeScript RT Reagent Kit with gDNA Eraser (Perfect Real Time; Takara, Dalian, China). For the target genes, qPCR experiments were performed in 72-well PCR plates using SYBR*® Premix Ex Taq*^TM^II-Tli RNaseH Plus (Takara, Dalian, China) in a 20-μL final reaction volume. PCR mixtures were carried out in molecular biology-grade water in the presence of 1×SYBR*® Premix*, 0.2 μM of the forward and reverse primers, and one to ten nanograms of cDNA template. Amplifications were performed using a Rotor-Gene Q real-time PCR detection system (Qiagen, Germany) with the following cycling conditions: 95 °C for 30 s, followed by [95 °C for 5 s; 60 °C for 20 s; 72 °C for 12 s] × 40 cycles and a melting curve analysis. 16S rRNA was selected as the reference gene, and the primers used were 338F/518R (Table 3). Untranscribed RNA was used as a control to ensure that no remaining genomic DNA could be detected. Reaction mixtures with water as the template were set as the negative control. Melting curve analysis with temperatures ranging from 72 ℃ to 95 ℃ at the end of the amplification cycles and agarose gel analyses of the amplification products were used to confirm the specificity of the qPCR products. All experiments were performed with four biological replicates, and each sample had four technical replicates. In addition, the mean estimated PCR efficiency for each amplicon group was calculated by the LinRegPCR program (version 2013.0) and used instead of the theoretical efficiency to increase the accuracy of the results^[43-45]^.

### Gene disruption and complementation

pEX18Tc-*hpdA* was constructed for gene knockout by fusing the erythromycin resistance gene and two upstream and downstream fragments of the target gene amplified with the primers shown in Table 3 to *Sac* I/*Hin*d III-digested pEX18Tc with the In-Fusion*®* HD Cloning Kit (TaKaRa, Dalian, China). The resulting plasmid pEX18Tc-*hpdA* was transformed into *E. coli* WM3064 (2,6-DAP auxotroph)-competent cells before conjugation with strain HP1 as described previously^[46]^. The double-crossover recombinants of HP1Δ*hpdA* were screened on LB plates containing 10% sucrose (w/v) and erythromycin. The obtained mutant strain HP1Δ*hpdA* was verified by PCR amplification and sequencing analysis (data not shown) using specific primers. Sequencing analysis was performed by YKang Biological Technology Co., Ltd. (Hangzhou, China). *orf9* and *orf10* were disrupted with the same procedure as *hpdA*.

pBBRBleo was constructed by replacing the kanamycin resistance gene in pBBR1MCS-2 with the bleomycin resistance gene. For gene complementation, pBBRBleo-*hpdA* was constructed by fusing the PCR products of *hpdA* amplified with the primers shown in Table 3 to *Kpn* I/*Xba* I-digested pBBRBleo. pBBRBleo-*hpdA* was used to transform *E. coli* WM3064 and then mated into the mutant strain HP1Δ*hpdA* by conjugation to obtain the complementary strain HP1Δ*hpdA*(pBBRBleo-*hpdA*). Moreover, pBBRBleo was also introduced into the mutant strain HP1Δ*hpdA* through conjugation to obtain the control strain HP1Δ*hpdA*(pBBRBleo).

### Growth and 3HP degradation analysis

The growth of and 3HP degradation by the wild-type strain HP1 and strains HP1Δ*hpdA*, HP1Δ*hpdA*(pBBRBleo-*hpdA*), and HP1Δ*hpdA*(pBBRBleo) were investigated. All four strains were cultivated in MSM with 1 g/L 3HP. Small samples of the medium were removed regularly for measurement of the strain growth and substrate concentration. The growth of strains was detected by measuring the optical density of the medium sample at 600 nm (OD_600_ nm) using a spectrophotometer. To determine the 3HP concentration in the medium, the cells were removed by centrifugation at 14,000×g for 2 min, and the supernatants were carefully collected after filtering through a 0.22 μm filter and used for high-performance liquid chromatography (HPLC) analysis. The growth of and 3HP degradation by other mutants were analyzed following the same methods as those for the mutant strain HP1Δ*hpdA*.

### Bioinformatics analysis

The domain annotation of *hpdA* was analyzed by BLAST analysis of NCBI using the Conserved Domain Database (CDD). The amino acid sequence encoded by *hpdA3* was analyzed by TMHMM (http://www.cbs.dtu.dk/services/TMHMM/), HMMTOP (http://www.enzim.hu/hmmtop/), TMpred (https://embnet.vital-it.ch/software/TMPRED_form.html) and SignalP-5.0 (http://www.cbs.dtu.dk/services/SignalP/) to predict the transmembrane domain (Tmd) and signal peptide (Sp).

### Heterologous expression of *hpdA*

To verify the function of *hpdA*, HpdA was heterologously expressed. *S. melonis* TY, *E. adhaerens* ZM04 (collection number, CCTCC AB 2019220) and *Alcaligenes faecalis* JQ135 were chosen as the expression hosts. The pRK415-*hpdA* series expression vector was generated with a fragment of *hpdA* just downstream of the promoter of the vector pRK415 (*Hin*d III/*Eco*R I-digested). After sequencing and obtaining the desired construction, the plasmid construct was transformed into *E. coli* WM3064 and then mated into *S. melonis* TY, *E. adhaerens* ZM04 and *A. faecalis* JQ135 to generate *S. melonis* TY-*hpdA*, *E. adhaerens* ZM04-*hpdA* and *A. faecalis* JQ135-*hpdA*, respectively. The expression of HpdA was induced by adding 0.1 mM IPTG. The cells were harvested by centrifugation at 6,000×g for 5 min and washed twice with 12 mM phosphate-buffered saline (PBS), pH 7.4, and the cell pellets were resuspended in MSM until an OD_600_ nm value of 1.0 was reached (resting cells). Then, the 3HP transformation ability of resting cells was determined. 3HP was added at a final concentration of 0.1 mg/mL in the resting cell suspension, which was then shaken at 200 rpm at 30 °C for 8 h. Then, samples were taken aseptically for UV and LC-MS analysis to detect the formation of the product.

### Construction, expression and cellular localization of GFP fusion protein

The full-length amino acid sequence of HpdA3 was fused with GFP at the N terminus. pGFPe*hpdA3* was constructed by fusing the PCR fragment of *hdpA3* to pGFPe (digested with *Eco*R I and *Bam*H I, yielding an 11-amino acid linker region (GSENLYFQGQF) followed by GFP). The primers used to construct the GFP fusion protein are listed in Table 3. The construct was transformed into *E. coli* BL21 (DE3) and grown in LB medium containing kanamycin. Transformant colonies verified by sequencing were grown in LB medium containing kanamycin at 37 ℃ to an OD_600_ nm of 0.5. Cultures were induced with 0.5 mM IPTG for 4 h at 30 ℃ on a rotary shaker (200 rpm). The cells were harvested at 10,000×g for 1 min at 4 ℃, washed twice with PBS (pH 7.4) and resuspended in PBS (pH 7.4). Samples were imaged under a confocal microscope with a 488-nm excitation filter and a 520-nm emission filter.

### Analytical methods

The concentration of 3HP was determined by HPLC with diode array detection. The column used was an Eclipse XDB-C18 reverse-phase column (5 μm; 4.6 × 250 mm; Agilent, USA) at 35 °C. The mobile phase was 20% (v/v) methanol and 80% (v/v) 0.1% triethylamine at a flow rate of 1.0 mL min^−1^, and the detection wavelength was set at 254 nm. LC-MS analysis to determine the heterologous expression product of *hpdA* was performed on a liquid chromatograph (Agilent 1200, USA) equipped with an Eclipse XDB-C18 reverse-phase column (5 μm; 4.6 × 250 mm; Agilent, USA) and an LCQ Deca XP Max MS instrument (Thermo Finnigan) with an electrospray interface (Turbo Ion Spray). The iron spray voltage was set at 3,000 V. Nitrogen was used as the sheath gas (60 arb) and auxiliary gas (15 arb). The capillary temperature was set at 350 ℃, and the capillary voltage was set at 10 V. The mobile phase was 95% (v/v) of 0.1% (v/v) formic acid and 5% (v/v) of methanol at a flow rate of 1 mL/min, and 10 μL of sample was injected. The column temperature was set at 35 ℃, and the detection wavelength was 310 nm. Positive electrospray ionization with continuous full scanning from m/z 50 to 350 was performed. Samples for HPLC and LC-MS analysis were centrifuged at 14,000×g for 2 min, and the supernatant was filtered through a 0.22 μm filter prior to injection.

## Data availability

The 16S rRNA sequence of *E. adhaerens* HP1 has been deposited in the GenBank database under the accession number MN083303. The Whole Genome Shotgun project of *E. adhaerens* HP has been deposited at DDBJ/ENA/GenBank under accession VHKK00000000. The version described in this paper is version VHKK01000000.

## Acknowledgments

This work was financially supported by grants from the National Natural Science Foundation of China (Nos. 41721001, 41630637 and 31800089).

