## Supplemental table S1 and S2, Figure S1-S3 for "Catabolism of 3-hydroxypyridine by *Ensifer adhaerens* HP1: a novel four-component gene encoding 3-hydroxypyridine dehydrogenase HpdA catalyzes the first step of biodegradation"

**Supplemental tables**

**Table S1** Accession data of bacterial strains containing the *3hpd* gene cluster.

**Table S2** Sequence similarity and identity of corresponding gene showed in Table S1 to *3hpd* gene.

**Supplemental figures**

**Figure S1** Mass spectra of 2,5-DHP as determined by LC-MS analysis.

**Figure S2** Transcriptional analysis of three other candidate 3HP monooxygenase genes.

**Figure S3** Mass spectra of 2,5-DHP as determined by LC-MS analysis.

**Table S1** Accession data of bacterial strains containing the *3hpd* gene cluster.

| Kingdom | Phylum | Class | Order | Family | Genus+Species+Strains[#] | Types[$] | Genome accession numbers [※] | hpdA1[&] | hpdA2 | hpdA3 | hpdA4 | orf7 | orf8 | orf9 | orf10 | hpdB | hpdC | hpdD | hpdE |
|---|---|---|---|---|---|---|---|---|---|---|---|---|---|---|---|---|---|---|---|
| Bacteria | Actinobacteria | Actinobacteria | Corynebacteriales | Mycobacteriaceae | *Mycobacterium* sp. GA-2829 | XVIII | NZ_LQIT01000038 | AU194_RS21550 | AU194_RS21545 | AU194_RS21525 | AU194_RS21520 | AU194_RS21500 | AU194_RS21530 | AU194_RS21515 | AU194_RS21510 | AU194_RS21540 | AU194_RS21535 | AU194_RS21505 | AU194_RS21555 |
| | | | | | *Mycobacterium* sp. MS1601 | XVII | NZ_CP019420 | BVC93_RS12860 | BVC93_RS12855 | BVC93_RS12835 | BVC93_RS12830 | BVC93_RS12805 | BVC93_RS12840 | BVC93_RS12825 | BVC93_RS12820 | BVC93_RS12850 | BVC93_RS12845 | BVC93_RS12815 | BVC93_RS12865 |
| | | | | | *Mycolicibacterium smegmatis* MKD8 | XVIII | NZ_CP027541 | D806_RS10515 | D806_RS10520 | D806_RS10540 | D806_RS10545 | D806_RS10565 | D806_RS10535 | D806_RS10550 | D806_RS10555 | D806_RS10525 | D806_RS10530 | D806_RS10560 | D806_RS10510 |
| | | | Geodermatophilales | Geodermatophilaceae | *Blastococcus* sp. DSM 46838 | XVII | NZ_FOND01000033 | BM142_RS23570 | BM142_RS23565 | BM142_RS23545 | BM142_RS23540 | BM142_RS23515 | BM142_RS23550 | BM142_RS23535 | BM142_RS23530 | BM142_RS23560 | BM142_RS23555 | BM142_RS23525 | BM142_RS23575 |
| | | | | | *Geodermatophilus ruber* strain DSM 45317 | XVII | NZ_FOSW01000005 | BMZ50_RS10500 | BMZ50_RS10505 | BMZ50_RS10525 | BMZ50_RS10530 | BMZ50_RS10555 | BMZ50_RS10520 | BMZ50_RS10535 | BMZ50_RS10540 | BMZ50_RS10510 | BMZ50_RS10515 | BMZ50_RS10545 | BMZ50_RS10495 |
| | | | | | *Geodermatophilus sabuli* strain DSM 46844 | XIX | NZ_OBDO01000012, NZ_OBDO01000019, NZ_OBDO01000025 | CRP39_RS23515 | CRP39_RS23520 | CRP39_RS25960 | CRP39_RS25955 | CRP39_RS25930 | CRP39_RS25965 | CRP39_RS25950 | CRP39_RS25945 | CRP39_RS26075 | CRP39_RS25970 | CRP39_RS25940 | CRP39_RS23510 |
| | | | Pseudonocardiales | Pseudonocardiaceae | *Amycolatopsis acidiphila* strain JCM 30562 | XX | NZ_VJZA01000001 | FNH06_RS00605 | FNH06_RS00600 | FNH06_RS00580 | FNH06_RS00575 | / | FNH06_RS00585 | FNH06_RS00570 | FNH06_RS00565 | FNH06_RS00595 | FNH06_RS00590 | FNH06_RS00560 | FNH06_RS00610 |
| | | | | | *Pseudonocardia ammonioxydans* strain CGMCC 4.1877 | XVIII | NZ_FOUY01000105, NZ_FOUY01000096 | BM093_RS34035 | BM093_RS34030 | BM093_RS33680 | BM093_RS33685 | BM093_RS33705 | BM093_RS33675 | BM093_RS33690 | BM093_RS33695 | BM093_RS33665 | BM093_RS33670 | BM093_RS33700 | BM093_RS34040 |
| | | | | | *Pseudonocardia autotrophica* strain NRRL B-16064 | XVII | NZ_JNYD01000043, NZ_JNYD01000052 | OQ00_RS35215 | OQ00_RS35220 | OQ00_RS36350 | OQ00_RS36355 | OQ00_RS36380 | OQ00_RS36345 | OQ00_RS36360 | OQ00_RS36365 | OQ00_RS35225 | OQ00_RS36340 | OQ00_RS36370 | OQ00_RS35210 |
| | | | | | *Pseudonocardia kunmingensis* strain DSM 45301 | XVIII | NZ_VFPA01000001 | FB558_RS00710 | FB558_RS00705 | FB558_RS00685 | FB558_RS00680 | FB558_RS00660 | FB558_RS00690 | FB558_RS00675 | FB558_RS00670 | FB558_RS00700 | FB558_RS00695 | FB558_RS00665 | FB558_RS00715 |
| | | | | | *Pseudonocardia oroxyli* strain CGMCC 4.3143 | XVII | NZ_FNBE01000014, NZ_FNBE01000023 | BLS43_RS21215 | BLS43_RS21220 | BLS43_RS27250 | BLS43_RS27245 | BLS43_RS27220 | BLS43_RS27255 | BLS43_RS27240 | BLS43_RS27235 | BLS43_RS27265 | BLS43_RS27260 | BLS43_RS27230 | BLS43_RS21210 |
| | | | | | *Pseudonocardia* sp. MH-G8 | XVIII | NZ_NKYF01000048, NZ_NKYF01000056 | CFP66_RS45465 | CFP66_RS45460 | CFP66_RS46565 | CFP66_RS46570 | CFP66_RS46590 | CFP66_RS46560 | CFP66_RS46575 | CFP66_RS46580 | CFP66_RS46550 | CFP66_RS46555 | CFP66_RS46585 | CFP66_RS45470 |
| Bacteria | | Rubrobacteria | Gaiellales | Gaiellaceae | *Gaiella occulta* strain F2-233 | XXI | NZ_QQZY01000003 | Gocc_RS08190 | Gocc_RS08195 | Gocc_RS08185 | Gocc_RS08200 | Gocc_RS08235 | Gocc_RS08180 | Gocc_RS08205 | Gocc_RS08215 | Gocc_RS08220 | Gocc_RS08175 | Gocc_RS08225 | Gocc_RS08210 |
| Bacteria | | Thermoleophilia | Solirubrobacterales | unclassified Solirubrobacterales | *Solirubrobacterales* bacterium 70-9 SCNpilot | XXII | MKSH01000023 | BGO11_17690 | BGO11_17695 | BGO11_17685 | BGO11_17700 | BGO11_17730 | / | BGO11_17705 | BGO11_17660 | BGO11_17710 | BGO11_17715 | BGO11_17725 | BGO11_17680 |
| Bacteria | Proteobacteria | Alphaproteobacteria | Rhizobiales | Hyphomicrobiaceae | *Hyphomicrobium* sp. 99 | II | NZ_KQ031382 | G359_RS05965 | G359_RS05970 | G359_RS05990 | G359_RS05995 | G359_RS06025 | G359_RS05960 | G359_RS06000 | G359_RS06005 | G359_RS05975 | G359_RS05980 | G359_RS06010 | G359_RS05985 |
| | | | | Rhizobiaceae | ***Ensifer adhaerens* HP1** | I | VHKK00000000 | | | | | | | | | | | | |
| | | | | unclassified Rhizobiales | *Rhizobiales* bacterium isolate AFS066724 | III | NZ_UCDB01000020, NZ_UCDB01000016 | DUN04_RS18740 | DUN04_RS18745 | DUN04_RS18750 | DUN04_RS18755 | DUN04_RS18730 | DUN04_RS18735 | DUN04_RS18760 | DUN04_RS18765 | DUN04_RS20775 | DUN04_RS20770 | DUN04_RS20780 | DUN04_RS20765 |
| | | | | Xanthobacteraceae | *Starkeya novella* isolate S2 | IV | QFQD01000024 | DI549_09330 | DI549_09325 | DI549_09320 | DI549_09315 | DI549_09270 | DI549_09335 | DI549_09310 | DI549_09305 | DI549_09340 | DI549_09345 | DI549_09300 | DI549_09350 |
| | | | Rhodobacterales | Rhodobacteraceae | *Confluentimicrobium* sp. EMB200-NS6 | V | NZ_CP010869 | TQ29_RS12750 | TQ29_RS12755 | TQ29_RS12775 | TQ29_RS12780 | TQ29_RS12800 | TQ29_RS12745 | TQ29_RS12785 | TQ29_RS12790 | TQ29_RS12760 | TQ29_RS12765 | TQ29_RS12795 | TQ29_RS12770 |
| Bacteria | | Betaproteobacteria | Burkholderiales | Alcaligenaceae | *Pusillimonas caeni* strain KCTC 42353 | VI | NZ_PDUW01000002 | CSC67_RS04575 | CSC67_RS04580 | CSC67_RS04585 | CSC67_RS04590 | CSC67_RS03835 | CSC67_RS04570 | CSC67_RS04595 | CSC67_RS04600 | CSC67_RS03810 | CSC67_RS03815 | CSC67_RS03805 | CSC67_RS04565 |

| Kingdom | Phylum | Class | Order | Family | Genus+Species+Strains[#] | Types[$] | Genome accession numbers [※] | hpdA1[&] | hpdA2 | hpdA3 | hpdA4 | orf7 | orf8 | orf9 | orf10 | hpdB | hpdC | hpdD | hpdE |
|---|---|---|---|---|---|---|---|---|---|---|---|---|---|---|---|---|---|---|---|
| | | | | | *Pusillimonas noertemannii* BS8 | VII | NZ_AMZF01000007, NZ_AMZF01000020, NZ_AMZF01000064 | ON18_RS08065 | ON18_RS08070 | ON18_RS08075 | ON18_RS08080 | ON18_RS15165 | ON18_RS08060 | ON18_RS08085 | ON18_RS08090 | ON18_RS02355 | ON18_RS15185 | ON18_RS02360 | ON18_RS08055 |
| | | | | | *Pusillimonas* sp. 17-4A | VIII | NZ_NQOU01000002, NZ_NQOU01000004 | CJO09_RS08040 | CJO09_RS08045 | CJO09_RS08050 | CJO09_RS08055 | / | CJO09_RS08035 | CJO09_RS08060 | CJO09_RS08065 | CJO09_RS08445 | CJO09_RS11500 | CJO09_RS08480 | CJO09_RS08340 |
| | | | | | *Pusillimonas* sp. EA3 | VIII | NZ_QPJH01000001, NZ_QPJH01000003 | DER48_RS00645 | DER48_RS00640 | DER48_RS00635 | DER48_RS00630 | / | DER48_RS00650 | DER48_RS00625 | DER48_RS00620 | DER48_RS00215 | DER48_RS05995 | DER48_RS00105 | DER48_RS00345 |
| | | | | | *Pusillimonas* sp. L52-1-41 | IX | NZ_NQYH01000001, NZ_NQYH01000003 | CJP73_RS05765 | CJP73_RS05760 | CJP73_RS05755 | CJP73_RS05750 | / | CJP73_RS05770 | CJP73_RS05745 | CJP73_RS05740 | CJP73_RS03350 | CJP73_RS03355 | CJP73_RS03345 | CJP73_RS05465 |
| | | | | | *Pusillimonas* sp. isolate EAC49 | X | NZQZ01000057, NZQZ01000060 | CML16_13435 | CML16_13430 | CML16_13425 | CML16_13420 | / | CML16_13440 | CML16_13415 | CML16_13410 | CML16_16430 | CML16_16435 | CML16_16425 | CML16_13135 |
| | | | | | *Pusillimonas* sp. isolate SAT20 | XI | PARW01000036, PARW01000050 | CML19_05175 | CML19_05170 | CML19_05165 | CML19_05160 | / | CML19_05180 | CML19_05155 | CML19_05150 | CML19_07850 | CML19_07855 | CML19_07845 | CML19_04875 |
| | | | | | *Pusillimonas* sp. isolate SAT110 | XI | PAYE01000002, PAYE01000024 | CML18_09675 | CML18_09680 | CML18_09685 | CML18_09690 | / | CML18_09670 | CML18_09695 | CML18_09700 | CML18_00450 | CML18_00445 | CML18_00455 | CML18_09975 |
| | | | | Comamonadaceae | *Acidovorax* sp. KKS102 | XII | NC_018708 | C380_RS17035 | C380_RS17040 | C380_RS17045 | C380_RS17050 | C380_RS17025 | C380_RS17030 | C380_RS17055 | C380_RS17060 | C380_RS17100 | C380_RS17095 | C380_RS17105 | C380_RS17090 |
| | | | | | *Comamonas testosteroni* I2 | XII | NZ_JMRS01000007 | Y901_RS04915 | Y901_RS04920 | Y901_RS04925 | Y901_RS04930 | Y901_RS04905 | Y901_RS04910 | Y901_RS04935 | Y901_RS04940 | Y901_RS04980 | Y901_RS04975 | Y901_RS04985 | Y901_RS04970 |
| | | | | | *Comamonas testosteroni* NBRC 100989 | XII | NZ_BAEC01000038 | COMTE_RS06070 | COMTE_RS06075 | COMTE_RS06080 | COMTE_RS06085 | COMTE_RS06060 | COMTE_RS06065 | COMTE_RS06090 | COMTE_RS06095 | COMTE_RS06140 | COMTE_RS06135 | COMTE_RS06145 | COMTE_RS06130 |
| | | | | | *Comamonas thiooxydans* strain S44 | XII | ADVQ01000052 | CTS44_14343 | CTS44_14348 | CTS44_14353 | CTS44_14358 | CTS44_14333 | CTS44_14338 | CTS44_14363 | CTS44_14368 | CTS44_14413 | CTS44_14408 | CTS44_14418 | CTS44_14403 |
| | | | | | *Comamonas* sp. A23 | XII | NZ_RSDT01000024 | EIC84_RS25190 | EIC84_RS25195 | EIC84_RS25200 | EIC84_RS25205 | EIC84_RS25180 | EIC84_RS25185 | EIC84_RS25210 | EIC84_RS25215 | EIC84_RS25260 | EIC84_RS25255 | EIC84_RS25265 | EIC84_RS25250 |
| | | | | | *Comamonas* sp. Z1 | XII | NZ_VOQT01000029 | FSY45_RS25890 | FSY45_RS25885 | FSY45_RS25880 | FSY45_RS25875 | FSY45_RS25900 | FSY45_RS25895 | FSY45_RS25870 | FSY45_RS25865 | FSY45_RS25820 | FSY45_RS25825 | FSY45_RS25815 | FSY45_RS25830 |
| | | | | | *Variovorax paradoxus* isolate S2 | XII | QFPP01000002 | DI563_00515 | DI563_00520 | DI563_00525 | DI563_00530 | DI563_00505 | DI563_00510 | DI563_00535 | DI563_00540 | DI563_00585 | DI563_00580 | DI563_00590 | DI563_00575 |
| Bacteria | | Gammaproteobacteria | Oceanospirillales | Halomonadaceae | *Halomonas* sp. MES3-P3E | XIII | NZ_PJBT01000014 | CXF87_RS04010 | CXF87_RS04005 | CXF87_RS04000 | CXF87_RS03995 | CXF87_RS03980 | CXF87_RS04015 | CXF87_RS03990 | CXF87_RS03985 | CXF87_RS03945 | CXF87_RS04020 | CXF87_RS04030 | CXF87_RS04025 |
| | | | | | *Salinicola peritrichatus* strain JCM 18795 | XVI | NZ_PZJU01000012 | DOM57_RS07165 | DOM57_RS07160 | DOM57_RS07155 | DOM57_RS07150 | DOM57_RS07130 | DOM57_RS07170 | DOM57_RS07145 | DOM57_RS07140 | DOM57_RS07080 | DOM57_RS07075 | DOM57_RS07085 | DOM57_RS07070 |
| | | | | | *Halomonadaceae* bacterium R4HLG17 | XIV | NZ_PVRV01000019 | C3Z11_RS02195 | C3Z11_RS02200 | C3Z11_RS02205 | C3Z11_RS02210 | C3Z11_RS02230 | C3Z11_RS02190 | C3Z11_RS02215 | C3Z11_RS02220 | C3Z11_RS02175 | C3Z11_RS02185 | C3Z11_RS02170 | C3Z11_RS02180 |
| | | | Xanthomonadales | Xanthomonadaceae | *Pseudoxanthomonas* sp. SGD-5-1 | XV | NZ_RRYM01000009, NZ_RRYM01000045 | EIN95_RS17490 | EIN95_RS17495 | EIN95_RS17500 | EIN95_RS17505 | / | EIN95_RS17485 | EIN95_RS17510 | EIN95_RS17515 | EIN95_RS15175 | EIN95_RS15180 | EIN95_RS15170 | EIN95_RS17480 |

Note:

1. # The strains were sorted alphabetically.
2. $ The organization types of the *3hpd* cluster were shown in Figure 9.
3. ※ All genome sequences can be found on NCBI website. Genome sequences were collected till to September 15, 2019.
4. & The locus_tag of the *3hpd* homologure genes.

| Kingdom | Phylum | Class | Order | Family | Genus+Species+Strains[#] | Types[$] | Genome accession numbers [※] | hpdA1[&] | hpdA2 | hpdA3 | hpdA4 | orf7 | orf8 | orf9 | orf10 | hpdB | hpdC | hpdD | hpdE |
|---|---|---|---|---|---|---|---|---|---|---|---|---|---|---|---|---|---|---|---|

5. / indicated that the corresponding homologure genes were not found in the draft genome sequence.

**Table S2** Sequence similarity and identity of corresponding gene showed in Table S1 to *3hpd* gene.

| Genus+Species+Strains[#] | *hpdA1* Identity (percent) | *hpdA2* Similarity (percent) | Identity (percent) | *hpdA3* Similarity (percent) | Identity (percent) | *hpdA4* Similarity (percent) | Identity (percent) | *orf7* Similarity (percent) | Identity (percent) | *orf8* Similarity (percent) | Identity (percent) | *orf9* Similarity (percent) | Identity (percent) | *orf10* Similarity (percent) | Identity (percent) | *hpdB* Similarity (percent) | Identity (percent) | *hpdC* Similarity (percent) | Identity (percent) | *hpdD* Similarity (percent) | Identity (percent) | *hpdE* Similarity (percent) | Identity (percent) | Similarity (percent) |
|---|---|---|---|---|---|---|---|---|---|---|---|---|---|---|---|---|---|---|---|---|---|---|---|---|
| *Mycobacterium* sp. GA-2829 | 64 | 78 | 65 | 76 | 42 | 60 | 50 | 66 | 44 | 63 | 36 | 52 | 54 | 67 | 65 | 77 | 53 | 65 | 45 | 62 | 59 | 73 | 52 | 67 |
| *Mycobacterium* sp. MS1601 | 63 | 76 | 64 | 76 | 40 | 58 | 50 | 66 | 46 | 63 | 38 | 54 | 55 | 70 | 63 | 77 | 52 | 65 | 41 | 58 | 57 | 73 | 52 | 67 |
| *Mycolicibacterium smegmatis* MKD8 | 64 | 77 | 67 | 77 | 42 | 60 | 50 | 67 | 46 | 62 | 38 | 55 | 55 | 67 | 64 | 77 | 53 | 66 | 45 | 61 | 58 | 74 | 53 | 67 |
| *Blastococcus* sp. DSM 46838 | 63 | 77 | 70 | 82 | 46 | 62 | 55 | 70 | 46 | 66 | 40 | 55 | 56 | 70 | 66 | 79 | 52 | 65 | 43 | 57 | 59 | 75 | 52 | 67 |
| *Geodermatophilus ruber* strain DSM 45317 | 64 | 76 | 66 | 80 | 45 | 60 | 56 | 69 | 45 | 64 | 38 | 53 | 57 | 72 | 66 | 80 | 52 | 65 | 42 | 55 | 60 | 75 | 50 | 66 |
| *Geodermatophilus sabuli* strain DSM 46844 | 62 | 75 | 69 | 82 | 46 | 61 | 56 | 69 | 47 | 65 | 41 | 55 | 57 | 70 | 66 | 80 | 53 | 65 | 41 | 55 | 62 | 77 | 53 | 67 |
| *Amycolatopsis acidiphila* strain JCM 30562 | 63 | 76 | 70 | 80 | 40 | 53 | 55 | 70 | / | / | 37 | 50 | 57 | 70 | 66 | 79 | 53 | 66 | 41 | 56 | 67 | 78 | 50 | 65 |
| *Pseudonocardia ammonioxydans* strain CGMCC 4.1877 | 63 | 75 | 64 | 77 | 42 | 59 | 54 | 68 | 48 | 65 | 43 | 57 | 56 | 70 | 65 | 77 | incomplete[a] | incomplete | 39 | 55 | 62 | 73 | 50 | 63 |
| *Pseudonocardia autotrophica* strain NRRL B-16064 | 63 | 75 | 67 | 77 | 43 | 58 | 54 | 70 | 47 | 66 | 39 | 53 | 57 | 71 | 66 | 78 | incomplete | incomplete | 39 | 56 | 64 | 76 | 51 | 64 |
| *Pseudonocardia kunmingensis* strain DSM 45301 | 62 | 76 | 64 | 79 | 41 | 58 | 55 | 68 | 47 | 64 | 37 | 51 | 57 | 70 | 65 | 78 | 53 | 66 | 40 | 57 | 60 | 74 | 50 | 63 |
| *Pseudonocardia oroxyli* strain CGMCC 4.3143 | 64 | 76 | 67 | 77 | 41 | 58 | 54 | 69 | 47 | 65 | 43 | 57 | 56 | 71 | 65 | 78 | 53 | 65 | 40 | 55 | 65 | 75 | 51 | 65 |
| *Pseudonocardia* sp. MH-G8 | 62 | 76 | 64 | 79 | 41 | 58 | 55 | 68 | 47 | 64 | 37 | 51 | 57 | 70 | 65 | 78 | 52 | 66 | 40 | 57 | 60 | 73 | 50 | 63 |
| *Gaiella occulta* strain F2-233 | 59 | 74 | 63 | 76 | 39 | 62 | 50 | 67 | 35 | 50 | 38 | 51 | 53 | 65 | 62 | 78 | 43 | 58 | 45 | 56 | 54 | 68 | 48 | 63 |
| *Solirubrobacterales* bacterium 70-9 SCNpilot | 62 | 76 | 64 | 79 | 45 | 67 | 52 | 69 | 35 | 54 | / | / | 53 | 70 | 50 | 69 | 43 | 58 | 35 | 52 | 54 | 69 | 48 | 63 |
| *Hyphomicrobium* sp. 99 | 67 | 80 | 70 | 83 | 49 | 64 | 54 | 70 | 48 | 65 | 59 | 74 | 63 | 75 | 71 | 82 | 56 | 68 | 40 | 56 | 61 | 74 | 50 | 66 |
| ***Ensifer adhaerens* HP1** | | | | | | | | | | | | | | | | | | | | | | | | |
| *Rhizobiales* bacterium isolate AFS066724 | 98 | 99 | 95 | 97 | 97 | 97 | 99 | 99 | 97 | 98 | 97 | 98 | 98 | 99 | 99 | 99 | 71 | 77 | 69 | 82 | 83 | 90 | 79 | 89 |
| *Starkeya novella* isolate S2 | 72 | 84 | 75 | 86 | 51 | 65 | 55 | 71 | 63 | 76 | 64 | 76 | 65 | 78 | 78 | 87 | 70 | 77 | 68 | 82 | 62 | 77 | 80 | 89 |
| *Confluentimicrobium* sp. EMB200-NS6 | 75 | 86 | 71 | 84 | 53 | 65 | 68 | 79 | 65 | 79 | 68 | 80 | 72 | 83 | 78 | 89 | 59 | 71 | 35 | 56 | 67 | 80 | 52 | 66 |
| *Pusillimonas caeni* strain KCTC 42353 | 63 | 78 | 63 | 80 | 44 | 62 | 61 | 76 | 25 | 42 | 57 | 71 | 60 | 73 | 72 | 82 | 31 | 49 | 55 | 70 | 41 | 51 | 64 | 75 |
| *Pusillimonas noertemannii* BS8 | 64 | 78 | 63 | 78 | 46 | 61 | 61 | 78 | 26 | 43 | 58 | 71 | 61 | 73 | 71 | 83 | 34 | 51 | 54 | 69 | 38 | 53 | 64 | 76 |
| *Pusillimonas* sp. 17-4A | 65 | 80 | 59 | 78 | 46 | 62 | 57 | 73 | / | / | 55 | 68 | 62 | 74 | 71 | 83 | 30 | 47 | 55 | 69 | 34 | 48 | 64 | 77 |
| *Pusillimonas* sp. EA3 | 65 | 79 | 60 | 78 | 46 | 60 | 56 | 73 | / | / | 57 | 70 | 61 | 74 | 71 | 83 | 30 | 46 | 55 | 69 | 35 | 49 | 64 | 77 |
| *Pusillimonas* sp. L52-1-41 | 65 | 79 | 59 | 78 | 46 | 62 | 57 | 73 | / | / | 55 | 68 | 62 | 75 | 71 | 83 | 30 | 46 | 55 | 69 | 35 | 49 | 64 | 77 |
| *Pusillimonas* sp. isolate EAC49 | 63 | 78 | 61 | 77 | 50 | 62 | 56 | 73 | / | / | 57 | 70 | 62 | 74 | 70 | 83 | 29 | 45 | 54 | 69 | 41 | 50 | 64 | 77 |

| Genus+Species+Strains[#] | *hpdA1* | | *hpdA2* | | *hpdA3* | | *hpdA4* | | *orf7* | | *orf8* | | *orf9* | | *orf10* | | *hpdB* | | *hpdC* | | *hpdD* | | *hpdE* | |
|---|---|---|---|---|---|---|---|---|---|---|---|---|---|---|---|---|---|---|---|---|---|---|---|---|
| | Identity (percent) | Similarity (percent) | Identity (percent) | Similarity (percent) | Identity (percent) | Similarity (percent) | Identity (percent) | Similarity (percent) | Identity (percent) | Similarity (percent) | Identity (percent) | Similarity (percent) | Identity (percent) | Similarity (percent) | Identity (percent) | Similarity (percent) | Identity (percent) | Similarity (percent) | Identity (percent) | Similarity (percent) | Identity (percent) | Similarity (percent) | Identity (percent) | Similarity (percent) |
| *Pusillimonas* sp. isolate SAT20 | 63 | 78 | 61 | 77 | 50 | 62 | 56 | 73 | / | / | 57 | 70 | 62 | 74 | 70 | 83 | 31 | 49 | 54 | 69 | 41 | 50 | 64 | 77 |
| *Pusillimonas* sp. isolate SAT110 | 65 | 79 | 59 | 78 | 46 | 62 | 57 | 73 | / | / | 55 | 68 | 62 | 75 | 71 | 83 | 30 | 45 | 55 | 68 | 42 | 51 | 63 | 75 |
| *Acidovorax* sp. KKS102 | 65 | 78 | 71 | 84 | 41 | 60 | 53 | 67 | 41 | 58 | 57 | 69 | 60 | 75 | 69 | 82 | 33 | 48 | 54 | 69 | 39 | 54 | 71 | 80 |
| *Comamonas testosteroni* I2 | 65 | 78 | 71 | 83 | 42 | 60 | 53 | 67 | 42 | 59 | 57 | 69 | 60 | 75 | 69 | 83 | 33 | 48 | 54 | 69 | 38 | 54 | 71 | 80 |
| *Comamonas testosteroni* NBRC 100989 | 65 | 78 | 71 | 83 | 42 | 60 | 53 | 67 | 42 | 58 | 57 | 69 | 60 | 75 | 69 | 83 | 33 | 48 | 54 | 69 | 38 | 54 | 71 | 80 |
| *Comamonas thiooxydans* strain S44 | 65 | 78 | 69 | 83 | 41 | 60 | 53 | 67 | 41 | 57 | 57 | 69 | 61 | 75 | 69 | 82 | 33 | 48 | 54 | 69 | 38 | 54 | 71 | 80 |
| *Comamonas* sp. A23 | 65 | 78 | 71 | 84 | 41 | 60 | 53 | 67 | 41 | 57 | 57 | 69 | 61 | 75 | 69 | 82 | 33 | 48 | 54 | 69 | 38 | 54 | 71 | 80 |
| *Comamonas* sp. Z1 | 65 | 78 | 71 | 84 | 41 | 60 | 53 | 67 | 41 | 57 | 57 | 69 | 61 | 75 | 69 | 82 | 33 | 48 | 54 | 69 | 38 | 54 | 71 | 80 |
| *Variovorax paradoxus* isolate S2 | 65 | 78 | 73 | 84 | 41 | 61 | 54 | 67 | 43 | 58 | 57 | 70 | 60 | 75 | 68 | 83 | 33 | 48 | 54 | 69 | 39 | 54 | 71 | 80 |
| *Halomonas* sp. MES3-P3E | 67 | 79 | 72 | 85 | 47 | 63 | 54 | 68 | 51 | 64 | 57 | 69 | 60 | 73 | 69 | 82 | 66 | 75 | 51 | 64 | 44 | 59 | 69 | 80 |
| *Salinicola peritrichatus* strain JCM 18795 | 68 | 79 | 67 | 81 | 45 | 61 | 54 | 69 | 50 | 61 | 59 | 72 | 62 | 74 | 69 | 82 | 71 | 78 | 71 | 83 | 84 | 89 | 83 | 90 |
| *Halomonadaceae* bacterium R4HLG17 | 68 | 79 | 71 | 85 | 47 | 62 | 56 | 71 | 45 | 62 | 58 | 71 | 62 | 74 | 69 | 81 | 70 | 77 | 53 | 65 | 85 | 90 | 69 | 80 |
| *Pseudoxanthomonas* sp. SGD-5-1 | 64 | 78 | 65 | 79 | 47 | 64 | 60 | 75 | / | / | 58 | 70 | 60 | 73 | 71 | 83 | 30 | 50 | 54 | 69 | 39 | 49 | 63 | 75 |

Note 1. # The strains were sorted the same as in Table S1.

2. a indicates that the ORF of the gene was incomplete

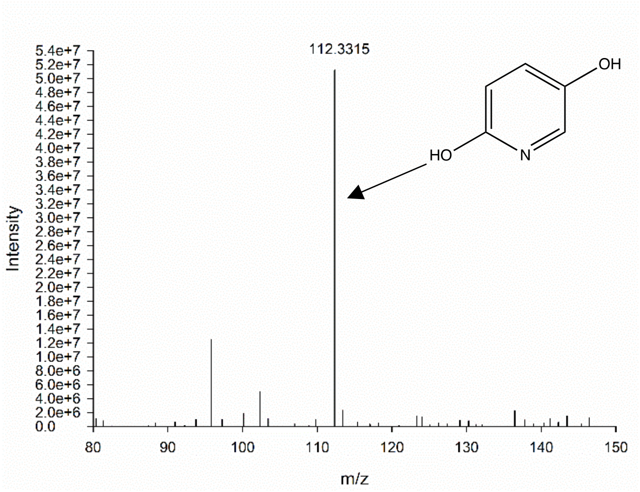


**Figure S1** Mass spectra of 2,5-DHP as determined by LC-MS analysis. A sample was taken from the strain HP1 grown with 3HP at 20 h (Figure 1).

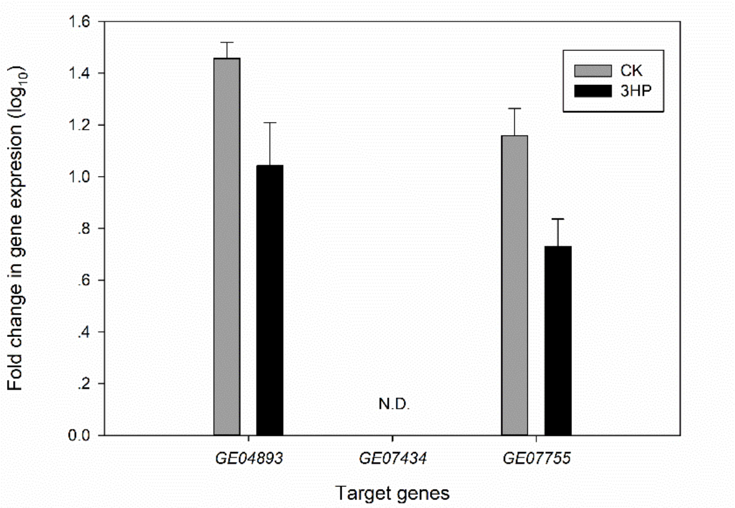


**Figure S2** Transcriptional analysis of three other candidate 3HP monooxygenase genes. RT-qPCR analysis of target gene transcripts produced in *E. adhaerens* HP1 grown with (black bars) or without (gray bars) 3HP. The expression levels of these three genes were normalized to the 16S rRNA expression level and are expressed as the fold change in expression in cells. The results presented in these histograms are the means of four independent experiments, and error bars indicate the standard error.

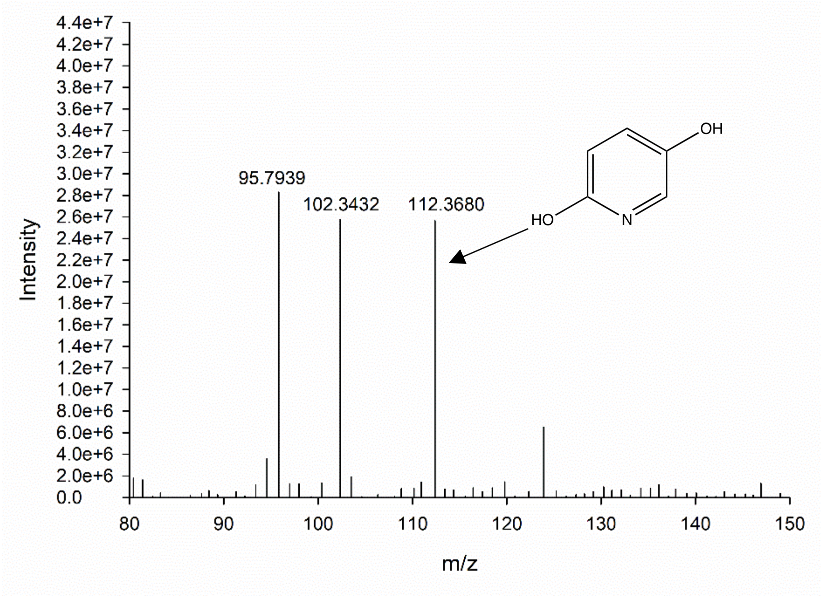


**Figure S3** Mass spectra of 2,5-DHP as determined by LC-MS analysis. A sample was taken from heterologous expression of *hphA* in strain ZM04 (Figure 6A).
